# Proteome coverage after simultaneous proteo-metabolome liquid-liquid extraction

**DOI:** 10.1101/2022.07.18.500507

**Authors:** A van Pijkeren, AS Egger, M Hotze, E Zimmermann, J Grander, A Gollowitzer, A Koeberle, R Bischoff, K Thedieck, M Kwiatkowski

## Abstract

Proteo-metabolomics is essential in systems biology and simultaneous proteo-metabolome extraction by liquid-liquid extraction (SPM-LLE) allows extraction of the metabolome and proteome from the same sample. Since the proteome is present as a pellet in SPM-LLE it must be solubilized for quantitative proteomics. Solubilization and proteome extraction is a critical factor in the information that can be obtained at the proteome level. In this study, we investigated the performance of two surfactants (sodium deoxycholate (SDC), sodium dodecyl sulfate (SDS)) and urea with respect to proteome coverage and extraction efficiency of an interphase proteome pellet generated by methanol-chloroform based SPM-LLE. We also investigated the extent to which the performance differs when the proteome is extracted from the interphase pellet or by direct cell lysis. Our study reveals that the proteome coverages between the two surfactants and urea for the SPM-LLE interphase pellet were very similar, but the extraction efficiencies differed significantly. While SDS led to enrichment of basic proteins, which were mainly ribosomal and ribonuclear proteins, urea was the most efficient extraction agent for simultaneous proteo-metabolome analysis. The results of our study also show that the performance of surfactants (SDC, SDS) for quantitative proteomics is better when the proteome was extracted by direct cell lysis and not from an interphase pellet. In contrast, the performance of urea for quantitative proteomics was significantly better when the proteome was extracted from an interphase pellet and by direct cell lysis.

## Introduction

Multiomics technologies are essential in modern life sciences and systems biology. However, integrating analyses across different omics platforms is still a major analytical challenge, especially with respect to sample preparation. A common assumption is that the various omics sample preparation techniques are platform dependent and mutually exclusive. This is intriguing, since classical methods, which have been used in lipid analysis for decades, such as the Bligh and Dyer ^1^ or the Folch ^2^ extractions, provide simultaneous access to the lipidome, the metabolome and the proteome. One reason for this discrepancy may be that sample preparation techniques used in metabolomics typically involve deproteinization steps to precipitate proteins by acid and/or organic solvents. Consequently, multiomics studies for which both the proteome and the metabolome need to be analyzed are often not done on the same sample, but rather two comparable samples are prepared independently, one for proteome and one for metabolome analysis ^3 4 5 6^. In recent years, methods for simultaneous proteo-metabolome extraction have been developed based on liquid-liquid extraction using methanol/chloroform ^7 8 9^ or methanol/methyl-*tert*-butyl-ether ^10^. In all these approaches, an interphase pellet containing the proteins is generated, and mostly urea is used to solubilize the proteins for subsequent proteome analysis. However, to date, there are no studies on the influence of different extraction agents and buffers on protein extraction efficiency from interphase pellets and consequently on proteome coverage in simultaneous proteo-metabolome liquid-liquid extraction (SPM-LLE) protocols. The choice of extraction agent is a critical step in multiomics analysis, as the chaotropes or surfactants used determine which part of the proteome is accessible for subsequent analysis.

There is also very limited information on whether and to what extent proteome coverages and extraction efficiencies differ between workup from an SPM-LLE interphase pellet and by direct cell lysis. Most studies focused on comparing the performance of chaotropic agents and surfactants on proteome coverage and digestion efficiency from proteome extracts generated by direct cell lysis ^11 12^.

Mass spectrometric based multiomics workflows are highly sophisticated multi-step experiments combining different methods, instruments and bioinformatics data processing workflows. The quality of multiomics experiments depends on the peculiarities and limitations of each step, with errors and/or biases of the individual steps propagating and accumulating throughout the experiment. Sample extraction is a critical step in the multiomics workflow, as chaotropes or surfactants determine which part of the proteome is available for subsequent analysis.

Here we describe a comparison of three different extraction buffer systems commonly used in proteomics (urea, sodium deoxycholate (SDC) and sodium dodecyl sulfate (SDS)) with respect to proteome coverage and extraction efficiency of an interphase proteome pellet generated by methanol-chloroform based SPM-LLE. We also investigated the extent to which the performance of each buffer system differs when the proteome is extracted from the interphase pellet or by direct cell lysis. For this study, we selected an immortalized human cell line (Human embryonic kidney 293T, HEK293T) as starting material and a label-free proteomics approach. Performance of the three proteome extraction buffer systems for SPM-LLE as well as between SPM-LLE and direct proteome extraction from cells were compared based on qualitative and quantitative proteome coverage, digestion efficiency, physicochemical properties (e.g., size, charge characteristics, and hydrophobicity) of extracted proteins and biological function. The results of this study will assist researchers in their choice of buffer system for proteome extraction in a simultaneous proteo-metabolome analysis as well as in conventional proteomics according to the focus of the biological question and the relevant protein populations.

## Experimental Section

### Chemicals

The chemicals used in this study are listed in the supplemental material (Supplemental Experimental Section).

### Cell culture and SPM-LLE

One million HEK293T cells were seeded in 6 well plates and grown for 48 hours at 37°C and 5% CO2 atmosphere in high-glucose (c=4.5 g/L) DMEM with FBS. The cells were washed three times with ice-cold PBS solution (pH 7.4). For simultaneous proteo-metabolome liquid-liquid extraction (SPM-LLE), 500 µL ice-cold methanol (MeOH) were added to the cells together with 20 µL of the [U-^13^C]-labelled yeast extract and 1 µL of the lipid standard (1,2-dimyristoyl-*sn*-glycero-3-phosphocholine (DMPC, c= 0.2 mM), 1,2-dimyristoyl-*sn*-glycero-3-phosphorylethanolamine (DMPE, c= 0.2 mM), 1,2,3-tri-myristoyl-glycerol (TMG, c= 0.2 mM)), followed by 500 µL ice-cold water. Lysates were transferred into a reaction tube (Eppendorf low binding tube, 2 mL, Eppendorf, Hamburg, Germany), followed by addition of 500 µL ice-cold chloroform (CHCl3) and incubation for 20 min at 4°C and 500 rpm on a thermo-shaker. Afterwards, samples were centrifugated for 5 min at (4°C and 16,000 x g). The polar and the non-polar phase were transferred into two new and separate reaction vials (Eppendorf low binding tube, 1.5 mL, Eppendorf, Hamburg, Germany), evaporated to dryness using an Eppendorf Concentrator Plus (Eppendorf, Hamburg, Germany) and stored at -80°C for further LC-MS analysis. The solid interphase pellet was evaporated to dryness using an Eppendorf Concentrator Plus (Eppendorf, Hamburg, Germany) and stored at -80°C for proteome extraction.

### Protein extraction from SPM-LLE interphase pellet using urea and tryptic digestion

The interphase pellets were dissolved in 60 µL urea buffer (8 M Urea, 100 mM ammonium bicarbonate (ABC), pH 8.3). The samples were diluted to a urea concentration of 2 M using 240 µL of 100 mM ABC (pH 8.3) and sonicated for 10 seconds at room temperature (Branson Ultrasonics™ Sonifier Modell 250 CE, Thermo Fisher Scientific, parameters: constant duty cycle, output control: 2). Total protein amount was quantified by Pierce Micro BCA Protein Assay Kit (Thermo Fisher Scientific, Dreieich, Germany) following the vendor protocol using a 1:50 dilution of a sample aliquot (V=10 µL) in HPLC-grade water (V=490 µL). For tryptic digestion, a volume containing 100 µg of total protein was transferred to a new reaction tube and made up to a final volume of 100 µL with 100 mM ABC (pH 8.3). For reduction, 1.05 µl of a dithiothreitol containing reduction buffer (1 M DTT, dissolved in 100 mM triethylammonium bicarbonate (TEAB), pH 8.3) was added and samples were incubated for 30 min at 55°C and 800 rpm on a thermo-shaker. For alkylation, 4.6 µL of iodoacetamide (IAA) containing alkylation buffer (0.5 M IAA, dissolved in 100 mM TEAB, pH 8.3) were added and samples were incubated for 30 min in the dark, followed by addition of 1.2 µL of reduction buffer to quench the alkylation reaction. Afterwards, 102.2 µL of 100 mM ABC was added and proteins were digested for 16 hours at 37°C using 5 µg of trypsin (dissolved in trypsin resuspension buffer, Promega, Walldorf, Germany). Tryptic digestion was stopped by addition of 2.5 µL 100% formic acid. The samples were centrifuged for 5 min (16,000 x g, RT), and the supernatants were used for reversed phase solid phase extraction.

### Protein extraction from SPM-LLE interphase pellet using sodium deoxycholate (SDC) and tryptic digestion

The interphase pellets were dissolved in 300 µL SDC buffer (2% w/v SDC, 100 mM TEAB, pH 8.3). The samples were heated for 5 min at 98°C, followed by sonication at room temperature (Branson Ultrasonics™ Sonifier Modell 250 CE, Thermo Fisher Scientific, parameters: 1x 10 seconds, constant duty cycle, output control: 2). Total protein amount was quantified by Pierce Micro BCA Protein Assay Kit (Thermo Fisher Scientific, Dreieich, Germany) following the vendor protocol using a 1:50 dilution of a sample aliquot (V=10 µL) in HPLC-grade water (V=490 µL). For tryptic digestion, a volume containing 100 µg of total protein was transferred to a new reaction tube and made up to a final volume of 100 µL with 100 mM TEAB (pH 8.3). Reduction, alkylation and quenching was performed as described above (see: Protein extraction from interphases of SPM-LLE using urea and tryptic digestion). Afterwards, 102.2 µL of 100 mM TEAB was added and proteins were digested for 16 hours at 37°C using 5 µg of trypsin (dissolved in trypsin resuspension buffer, Promega, Walldorf, Germany). Tryptic digestion was stopped by addition of 2.5 µL 100% formic acid. The samples were centrifuged for 5 min (16,000 x g, RT) to remove the precipitated SDC. The supernatants were used for reversed phase solid phase extraction.

### Protein extraction from SPM-LLE interphase pellet using sodium dodecyl sulfate (SDS) and tryptic digestion

The interphase pellets were dissolved in 300 µL sodium dodecyl sulfate (SDS) buffer (1 % SDS, 100 mM ABC, pH 8.3). The samples were heated for 5 min at 98°C, followed by sonication at room temperature (Branson Ultrasonics™ Sonifier Modell 250 CE, Thermo Fisher Scientific, parameters: 1x 10 seconds, constant duty cycle, output control: 2), and total protein amount was quantified with a Pierce Micro BCA Protein Assay Kit (Thermo Fisher Scientific, Dreieich, Germany) following the vendor instruction using a 1:50 dilution of a sample aliquot (V=10 µL) in HPLC-grade water (V=490 µL). 100 µg of total protein was used for tryptic digestion using single-pot solid-phase-enhanced sample preparation (SP3) procedure ^13^. For more details, see the supplemental material (Supplemental Experimental Section). The supernatants of SP3 procedure that contain the tryptic peptides were used for reversed phase solid phase extraction.

### Protein extraction by direct cell lysis using urea and tryptic digestion

Six well dishes were washed three times with 300 µL PBS. Cells were lysed by addition of 60 µL of urea containing buffer (8 M urea, 100 mM ABC, pH 8.3). The samples were diluted to a final urea concentration of 2 M using 240 µL of 100 mM ABC (pH 8.3) and sonicated for 10 seconds at room temperature (Branson Ultrasonics™ Sonifier Modell 250 CE, Thermo Fisher Scientific, parameters: constant duty cycle, output control: 2). Quantification of total protein amounts using Pierce Micro BCA Protein Assay Kit (Thermo Fisher Scientific, Dreieich, Germany) and tryptic digestion were performed as described above (see: Protein extraction from interphases of SPM-LLE using urea and tryptic digestion).

### Protein extraction by direct cell lysis using sodium deoxycholate (SDC) and tryptic digestion

Six well dishes were washed three times with 300 µL PBS. Cells were lysed by addition of 300 µL SDC buffer (2% w/v SDC, 100 mM TEAB, pH 8.3). The samples were heated for 5 min at 98°C, followed by sonication at room temperature (Branson Ultrasonics™ Sonifier Modell 250 CE, Thermo Fisher Scientific, parameters: 1x 10 seconds, constant duty cycle, output control: 2). Quantification of total protein amounts using Pierce Micro BCA Protein Assay Kit (Thermo Fisher Scientific, Dreieich, Germany) and tryptic digestion were performed as described above (see: Protein extraction from interphases of SPM-LLE using urea and tryptic digestion).

### Protein extraction by direct cell lysis using sodium dodecyl sulfate (SDS) and tryptic digestion

Six well dishes were washed three times with 300 µL PBS. Cells were lysed by addition of 300 µL SDS buffer 1 % SDS, 100 mM ABC, pH 8.3). The samples were heated for 5 min at 98°C, followed by sonication at room temperature (Branson Ultrasonics™ Sonifier Modell 250 CE, Thermo Fisher Scientific, parameters: 1x 10 seconds, constant duty cycle, output control: 2). Quantification of total protein amounts using Pierce Micro BCA Protein Assay Kit (Thermo Fisher Scientific, Dreieich, Germany), single-pot solid-phase-enhanced sample preparation (SP3) and tryptic digestion were performed as described above (see: Protein extraction from SPM-LLE interphase pellet using sodium dodecyl sulfate (SDS) and tryptic digestion).

### Reversed phase solid phase extraction (RP-SPE)

Samples were purified by RP-SPE prior to LC-MS analysis using OASIS HLB cartridges (Oasis HLB, 1 cc Vac Cartridge, 30 mg Sorbent, Waters, Manchester, UK) and a pressure manifold (Waters SPE Manifold, Waters, Manchester, UK). For more details, see the supplemental material (Supplemental Experimental Section).

### Proteome analysis by LC-MS/MS

Dried peptide samples were dissolved in 80 µL of 0.1% FA, and 1 µL of the samples were injected into a nano-ultra pressure liquid chromatography system (Dionex UltiMate 3000 RSLCnano pro flow, Thermo Scientific, Bremen, Germany) coupled via electrospray-ionization (ESI) to a tribrid orbitrap mass spectrometer (Orbitrap Fusion Lumos, Thermo Scientific, San Jose, CA, USA). The samples were loaded (15 µL/min) on a trapping column (nanoE MZ Sym C18, 5 μm, 180 µm x 20mm, Waters, Germany, buffer A: 0.1 % FA in HPLC-H2O; buffer B: 80 % ACN, 0.1 % FA in HPLC-H2O) with 5% buffer B. After sample loading the trapping column was washed for 2 min with 5% buffer B (15 μL/min) and the peptides were eluted (250 nL/min) onto the separation column (nanoEase MZ PST CSH, 130 A, C18 1.7 μm, 75 μm x 250mm, Waters, Germany; buffer A: 0.1 % FA in HPLC-H2O; buffer B: 80 % ACN, 0.1 % FA in HPLC-H2O). The peptides were separated using a total gradient of 110 min (5% B to 37.5% B in 90 min, 37.5% B to 62.5% B in 25 min) and analyzed in data dependent acquisition mode using the orbitrap for MS1 scans and the ion trap for MS2 scans. For more details, see the supplemental material (Supplemental Experimental Section).

### Bioinformatics data processing of proteome LC-MS/MS data

LC-MS/MS raw data were processed and quantified with MaxQuant (version 1.6.5.0). Peptide and protein identification were carried out with Andromeda. LC-MS/MS data was searched a human database (SwissProt, 20,431 entries, downloaded 19.08.2019, https://www.uniprot.org/) and a contaminant database (239 entries). For database search, a precursor mass tolerance of 6 ppm and a fragment mass tolerance of 0.5 Da were used. For peptide identification, two missed cleavages were allowed, a carbamidomethylation of cysteines was used as a static modification, and oxidation of methionine residues and acetylation of protein N-termini were allowed as variable modifications. Peptides and proteins were identified with an FDR of 1%. Proteins were quantified with the MaxLFQ algorithm considering only unique peptides, a minimum ratio count of two unique peptide and match between runs. The post-processing of the data was performed in R (version 4.0.3) and RStudio (version 1.4.1106). For more details, see the supplemental material (Supplemental Experimental Section).

### Analysis of polar metabolites and lipids by Ion Chromatography-Single Ion Monitoring-Mass Spectrometry (IC-SIM-MS) and multiple reaction monitoring mass spectrometry (LC-MRM-MS)

Polar metabolites and lipids were analyzed and quantified by IC-SIM-MS and LC-MRM-MS ^14 15 16^, respectively. Details of IC-SIM-MS and LC-MRM-MS analysis and bioinformatics data processing are described in the supplemental experimental section.

## Results and Discussion

### Access to lipids and polar metabolites by liquid-liquid extraction with chloroform and methanol

We used a MeOH-CHCl3-based SPM-LLE to extract the polar metabolome (MeOH phase), non-polar metabolome including lipids (CHCl3 phase) and the proteome (interphase pellet). Although the main focus of this work was to evaluate the proteome accessibility of the interphase pellet, we also exemplarily quantified 12 lipids covering triglycerides (TAG) and phospholipids (PC, PI, LPE and TAG) as well as 19 polar metabolites. All lipids (intracellular), including the added internal standards, were quantified by LC-MRM-MS with a coefficient of variation (CV) below 15% (Figure S1). Results of the IC-SIM-MS analysis of polar metabolites showed that all internal standards were quantified with CV values below 15% (Figure S2). For endogenous, intracellular polar metabolites, 4 out of 19 had CV values above 20% (α-ketoglutarate (CV=27.6%), fructose-1,6-bisphosphate (CV=28.1%), glucose-1-phosphate (CV=24.2%), 6-phosphogluconate (CV=22.3%)), whereas most metabolites (10 of 19) were quantified with CV values below 15%. From 7 extracellular polar metabolites, 5 were quantified with CV values below 15%, while malate (CV=21.9%) and fumarate (CV=22.0%) had CV values that were slightly above 20%. These results confirmed that lipids and polar metabolites can be quantified by CHCl3-MeOH-based LLE.

### Total protein yield

To investigate proteome accessibility by MeOH-CHCl3-based SPM-LLE, we used buffers containing either urea (dissolved in 100 mM ammonium bicarbonate, pH 8.3), SDC (dissolved in 100 mM triethylammonium bicarbonate, pH 8.3) or SDS (dissolved in 100 mM ammonium bicarbonate, pH 8.3) to solubilize the interphase pellet. We first determined the total protein amount using a colorimetric bicinchoninic acid assay. The average number of cells subjected to SPM-LLE was 4.3 × 10^6^ cells. Solubilization with SDS and urea exhibited little variation while solubilization with SDC was more variable. Usage of SDS provided the highest protein yield (1332 µg ± 44 µg) followed by SDC (784 µg ± 175 µg) and urea (659 µg± 49 µg) (Figure 1A). This shows that SDS is the most efficient solubilizing agent for proteins from the interphase pellet.

**Figure 1:**
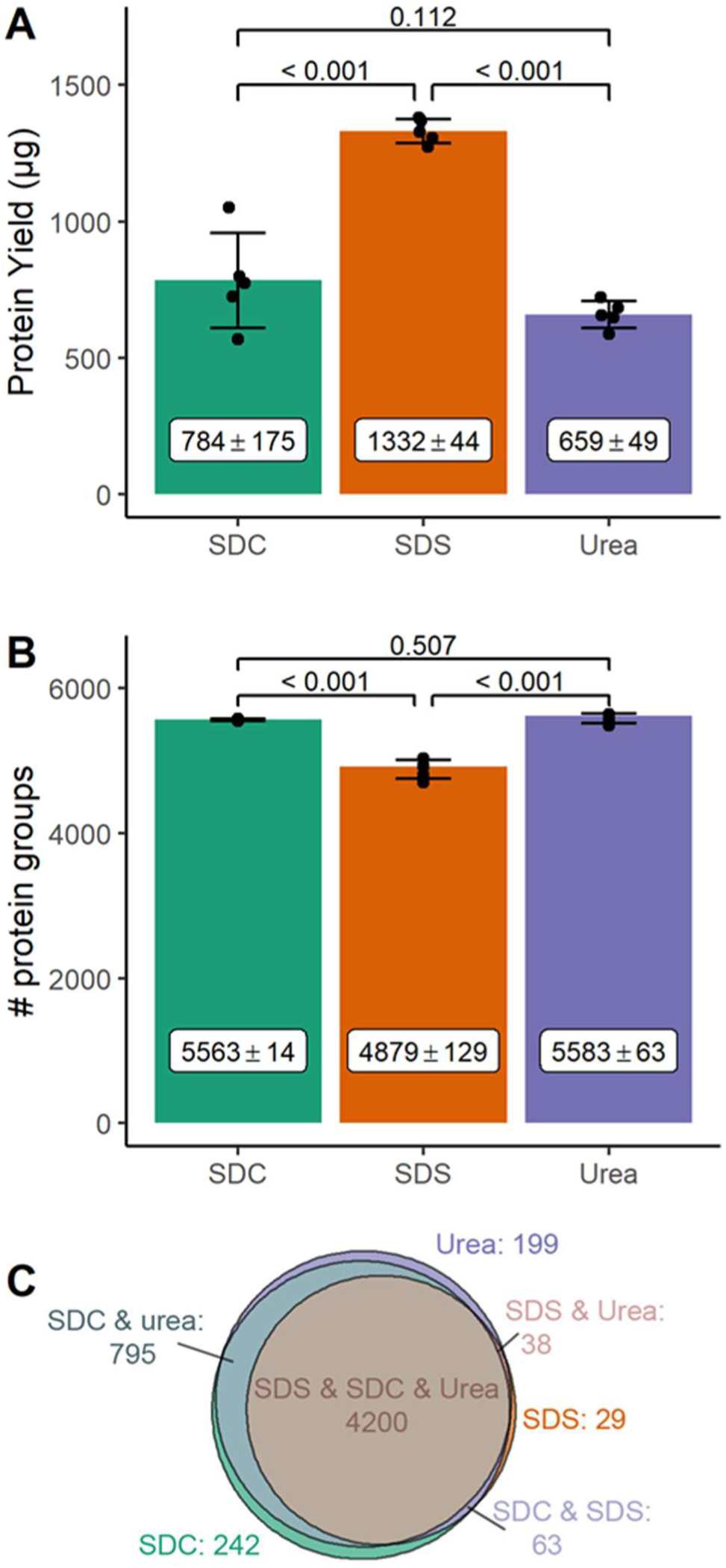
Protein yield (A) and number of proteins identified in the solubilized interphase pellets (B, C). A) Protein yield (µg) and B) number of identified proteins from the interphase pellets solubilized by sodium deoxycholate (SDC), sodium dodecyl sulfate (SDS), and urea. Bar graph: mean with standard deviation. Statistical analyses with two-tailed unpaired *t*-test. C) Number of proteins reproducibly identified in all independent experiments (n = 5, biological replicates) with the indicated buffer systems. Average number of cells: 4.3 × 10^6^ cells.

### Qualitative analysis of the solubilized proteome

To compare the composition of the proteome released with urea, SDS, or SDC, we used a single shot bottom-up label-free proteomics approach. Equal amounts of proteins extracted by urea, SDC or SDS were digested with trypsin. For the samples solubilized by SDS, the single-pot solid-phase-enhanced sample preparation (SP3) procedure was used to remove SDS prior to trypsin digestion ^13^. SDC was removed after tryptic digestion by acid precipitation and the urea concentration was reduced to less than 1 M prior to tryptic digestion by dilution. The resulting peptides were desalted by reversed phase solid phase extraction (RP-SPE) and dried. The dried peptide samples were dissolved in equal volumes and subjected to label free LC-MS/MS analysis.

Each solubilizing agent yielded reproducible numbers of identified proteins followed by SDC and SDS (Figure 1B). Urea (nurea= 5583 ± 63) and SDC (nSDC= 5563 ± 14) led to comparable numbers of identifications. In contrast, SDS (nSDS= 4879 ± 129) yielded a significantly lower number of identifications despite the better efficiency in solubilizing proteins from interphase pellets. The lower number of identified proteins suggests that the SP3 method is more prone to sample loss and/or introduces a bias towards certain proteins so that coverage is overall less during sample preparation compared to acid precipitation (SDC) or dilution (urea).

75.5% of all proteins (n= 4200) were reproducibly identified with all three solubilizing agents (Figure 1C, Table S1). The distributions of the identified proteins across the main GO cellular component categories (membrane proteins, nuclear proteins, cytoplasmatic proteins, Figure S3 A) and physicochemical properties (hydrophobicity, molecular weight, isoelectric point, Figure S3 B-D) were similar for the three different extraction systems. For proteins exclusively identified in urea, SDC or SDS (Figure S3 E-G), we observed no differences in hydrophobicity (Figure S3 E) or molecular weight (Figure S3 F). Proteins exclusively identified upon solubilization with SDS showed a shift towards higher, more basic isoelectric points (Figure S3 G).

We investigated the extent to which urea, SDC and SDS had an effect on missed cleavages (MC) during tryptic digestion. Tryptic digestion of the interphase pellet in presence of urea showed the lowest number of missed cleavages (MC0= 75%, MC1= 22%, MC2= 3%), followed by SDC (MC0= 64%, MC1= 29%, MC2= 7%) and SDS (MC0= 51%, MC1= 36%, MC2= 13%) (Figure S4, Table S2-S4).

In conclusion, from a qualitative perspective, urea, SDS and SDC-based proteome extractions from the SPM-LLE interphase pellet provide access to very similar proteomes, with SDS showing a tendency towards proteins with a higher isoelectric point. More obvious differences were observed for protein identification and tryptic digestion efficiency. SDS showed significantly lower number of identified proteins compared to urea and SDC, and the highest number of missed cleavages. Using urea resulted in the lowest number of missed cleavages and highest digestion efficiency compared to surfactants.

### Quantitative analysis of the solubilized proteome

To examine differences in quantitative access to the proteome, we performed an unsupervised multivariate analysis (principal component analysis, PCA). The five independent experiments of urea, SDC and SDS extractions formed clusters (Figure 2 A) that were clearly separate from each other in the first and second component, with the first component accounting for 64.1% of the summative variance and the second for 19.7%. Z-score hierarchical clustering based on squared Euclidean distance measure showed a similar result (Figure S 5).

**Figure 2:**
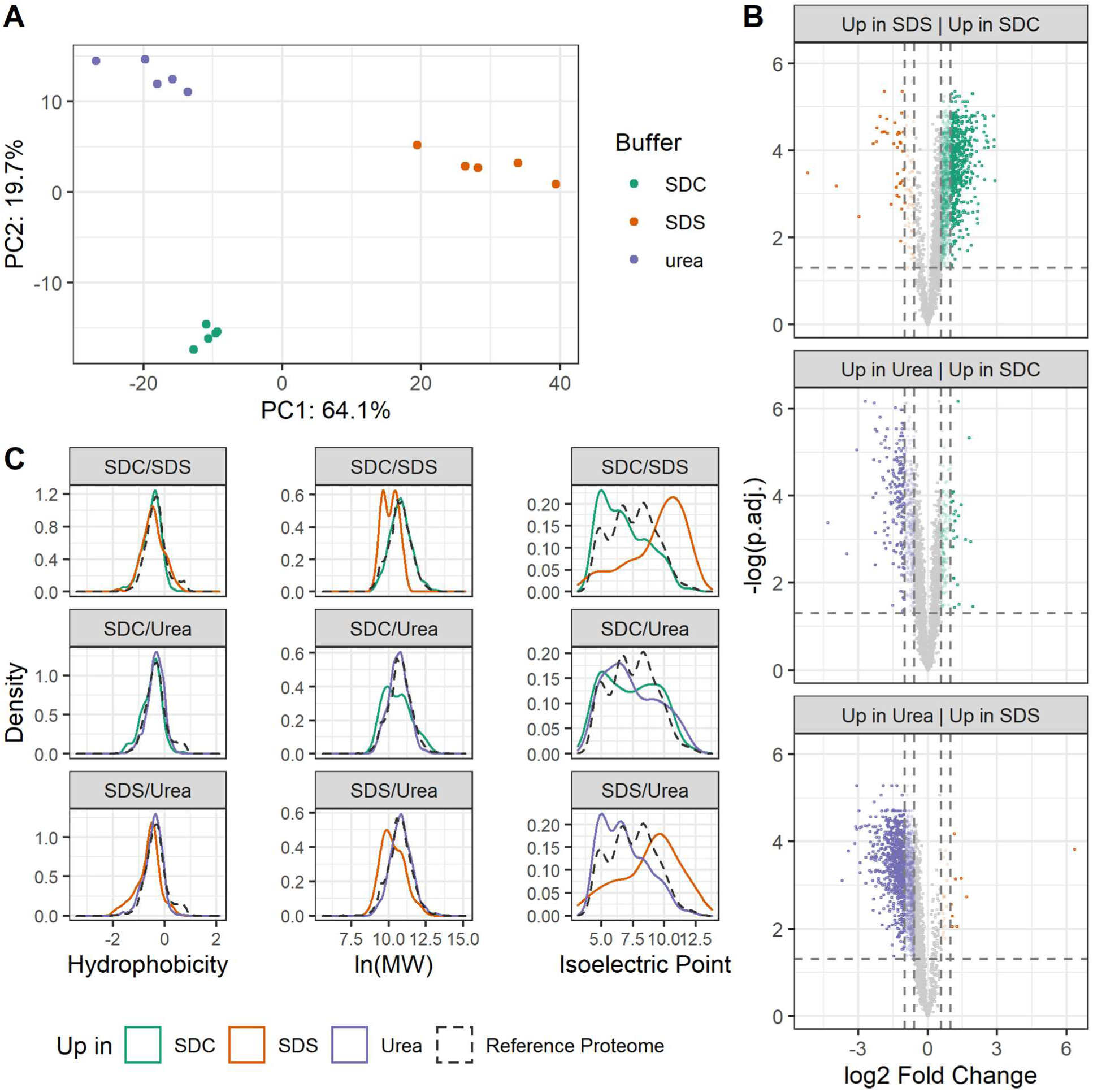
Quantitative analysis of proteins extracted from interphase pellets of simultaneous proteo-metabolomics liquid-liquid extractions (SPM-LLE). A) Principal component analysis of proteins extracted from the SPM-LLE interphase pellets using sodium deoxycholate (SDC, green), sodium dodecyl sulfate (SDS, orange), or urea (purple). Protein abundance levels, reproducibly quantified in all independent experiments and conditions, were used as input for PCA. B) Comparison of efficiency to extract proteomes from the SPM-LLE interphase pellet with SDC (green), SDS (orange), or urea (purple). Significance threshold for enrichment: adjusted *p*-value ≤ 0.05 (two-tailed unpaired *t*-test, Benjamini-Hochberg correction), fold change (FC) of 1.5: colored transparent dots, FC of ≥ 2: colored dots). C) Physicochemical properties of proteins enriched in the SPM-LLE interphase pellets (colored solid lines, FC ≥ 1.5, adjusted *p*-value ≤ 0.05) as compared to the human reference proteome (SwissProt, uniprot.org, black dashed line). n= 5 independent experiments (biological replicates).

To compare extraction efficiency, the differentially extracted proteins were visualized in volcano-plots (Figure 2 B). Proteins were considered significantly and differentially extracted at a threshold fold-change of at least 1.5 and an adjusted *p*-value below 0.05 (*t*-test, Benjamini-Hochberg correction). Based on these criteria, 1409 proteins were extracted more efficiently in SDC versus 91 in SDS. Comparing SDC with urea, 167 proteins were extracted more efficiently in SDC and 488 in urea. Comparing urea with SDS, 1596 proteins were extracted more efficiently in urea and 29 in SDS. These results show that urea provides the most efficient proteome extraction from the SPM-LLE interphase pellet.

While the differentially extracted proteomes did not exhibit differences in hydrophobicity (Figure 2 C), SDS extracted proteins of lower molecular weight more efficiently than SDC and urea. Likewise, SDC showed a trend to extract proteins with a lower molecular weight more efficiently than urea. In keeping with the qualitative analysis, proteins that were extracted more efficiently in SDS showed a shift to higher, more basic isoelectric points. GO enrichment analysis of cellular components (GO:CC) showed a significant enrichment of cytosolic as well as intracellular and membrane-bound organelle proteins in urea and SDC compared to SDS (Figure S6 A, F). In contrast, ribosomal and ribonuclear proteins were more efficiently enriched in SDS compared to urea and SDC (Figure S6 B, E). Our quantitative analysis showed that SDS-based extraction resulted in specific enrichment of basic ribosomal and ribonuclear proteins (Figure S5, Figure S7). Sixty-nine of the 92 enriched proteins have an isoelectric point greater than 9, which explains the observed shift to higher isoelectric points for proteins extracted more efficiently with SDS (Figure 2 C). While these ribosomal and ribonuclear proteins were underrepresented in the SDC proteome, this effect was less pronounced for the proteome extracted with urea (Figure S6 B, D, Figure S7). One possible explanation for this observation are the anionic properties of the sulfate group of SDS, which disrupts electrostatic interactions between positively charged proteins and the negatively charged RNA leading to a more efficient extraction of the ribosomal and ribonuclear proteins from the interphase pellet, which contains not only proteins but also nucleic acids.

In simultaneous proteo-metabolome analysis, the coverage of proteins related to metabolic pathways is of particular relevance for integration of the proteome and metabolome data. We therefore investigated the extraction efficiency for proteins involved in glycolysis and gluconeogenesis, the tricarboxylic acid (TCA) cycle, the pentose phosphate pathway (PPP), amino acid metabolism, and glycerolipid and glycerophospholipid metabolism. A qualitative comparison of the reproducibly identified proteins showed a similar coverage of all metabolic pathways with SDS, SDC, and urea (Figure S8). Quantitative analysis of extraction efficiency showed that proteins related to carbohydrate, lipid, and amino acid metabolism were best extracted from the interphase of SPM-LLE with urea (Table S5, Figure S7, Figure S9), whereas the lowest extraction efficiency was achieved with SDS. SDC showed higher efficiency than SDS in the extraction of proteins that are related to metabolic pathways.

### Proteome coverage of the SPM-LLE interphase pellet compared to extraction by direct cell lysis

We determined the extent to which proteome extraction from the interphase pellet by SPM-LLE introduces a systematic bias compared with the more commonly used proteome extraction by direct cell lysis ^17 18^. For direct cell lysis and proteome extraction (average number of cells: 4.3 × 10^6^ cells), SDS provided the highest protein yield (1002 µg ± 137 µg) followed by SDC (926 µg ± 66 µg) and urea (856 µg± 48 µg) (Figure S10 A). In contrast to solubilization of the interphase protein pellet, the use of SDS, SDC, and urea in direct protein extraction from cells showed no significant differences in total protein yield. For SDS, the total protein yield was lower with direct cell lysis than with extraction from the interphase protein pellet, but for SDC and urea, direct cell lysis resulted in higher protein yields (Figure 1 A, Figure S10 A).

For direct proteome extraction from cells, SDC yielded the highest number of identified proteins (nSDC= 5625 ± 34), followed by urea (nurea= 5577 ± 37) and SDS (nSDS= 5286 ± 66) (Figure S10 B). As for SPM-LLE, the use of SDS resulted in a significantly lower number of identified proteins compared to SDC and urea, although the difference was not as pronounced for direct proteome extraction from cells as for the interphase pellet (Figure 1B). 82.8% of proteins (n= 4646) were reproducibly identified using urea, SDC and SDS for direct cell lysis and proteome extraction (Figure S10 C, Table S6). When comparing the identified proteins, we did not find differences in proteome coverage between proteome extraction by direct cell lysis and SPM-LLE. This confirms results by Nakayasu et al. using urea for proteome extraction by direct cell lysis and interphase pellets after CHCl3-MeOH extraction ^9^. In our study, we observed a higher number of identified proteins by direct cell lysis for SDC and SDS, whereas the number of identified proteins for urea was higher in SPM-LLE interphase pellets (Figure S11 A-C). Most proteins were reproducibly identified in both by direct cell lysis and from SPM-LLE pellets when SDC (92.5%), SDS (88.5%), or urea (91.9%) were used (Figure S11 D-F).

However, we observed considerable differences in the efficiency of tryptic digestion between proteome extraction from SPM-LLE interphase pellets (Figure S4) and by direct cell lysis (Figure S12). Efficiency was improved by direct cell lysis for both SDC (Table S2) and SDS (Table S3), whereas urea showed better digestion efficiency using the SPM-LLE interphase pellet (Table S4).

PCA analysis showed that the individual experiments clustered together each and that all conditions (extraction agents, SPM-LLE interphase pellet, proteome extraction by direct cell lysis) were clearly separated from each other in the first (50.5%), second (20.1%) and third (11.5%) component (Figure 3A). Both SDC and SDS extracted proteins more efficiently by direct cell lysis (SDC: 305, SDS: 325) than from SPM-LLE interphase pellets (SDC: 118, SDS: 22) (Figure 3B; fold change threshold 1.5, adjusted p-value < 0.5, Benjamini-Hochberg correction). In contrast, urea-based protein extraction was more efficient from the SPM-LLE interphase pellet (136) as compared to direct cell lysis (103). Comparing the total number of proteins whose abundances differed significantly (adjusted p-value < 0.05, Benjamini-Hochberg correction) regardless of the fold-change threshold, our results show that a considerably larger number of proteins was extracted more efficiently from the SPM-LLE interphase pellet (n= 915) with urea than by direct cell lysis (n= 214).

**Figure 3:**
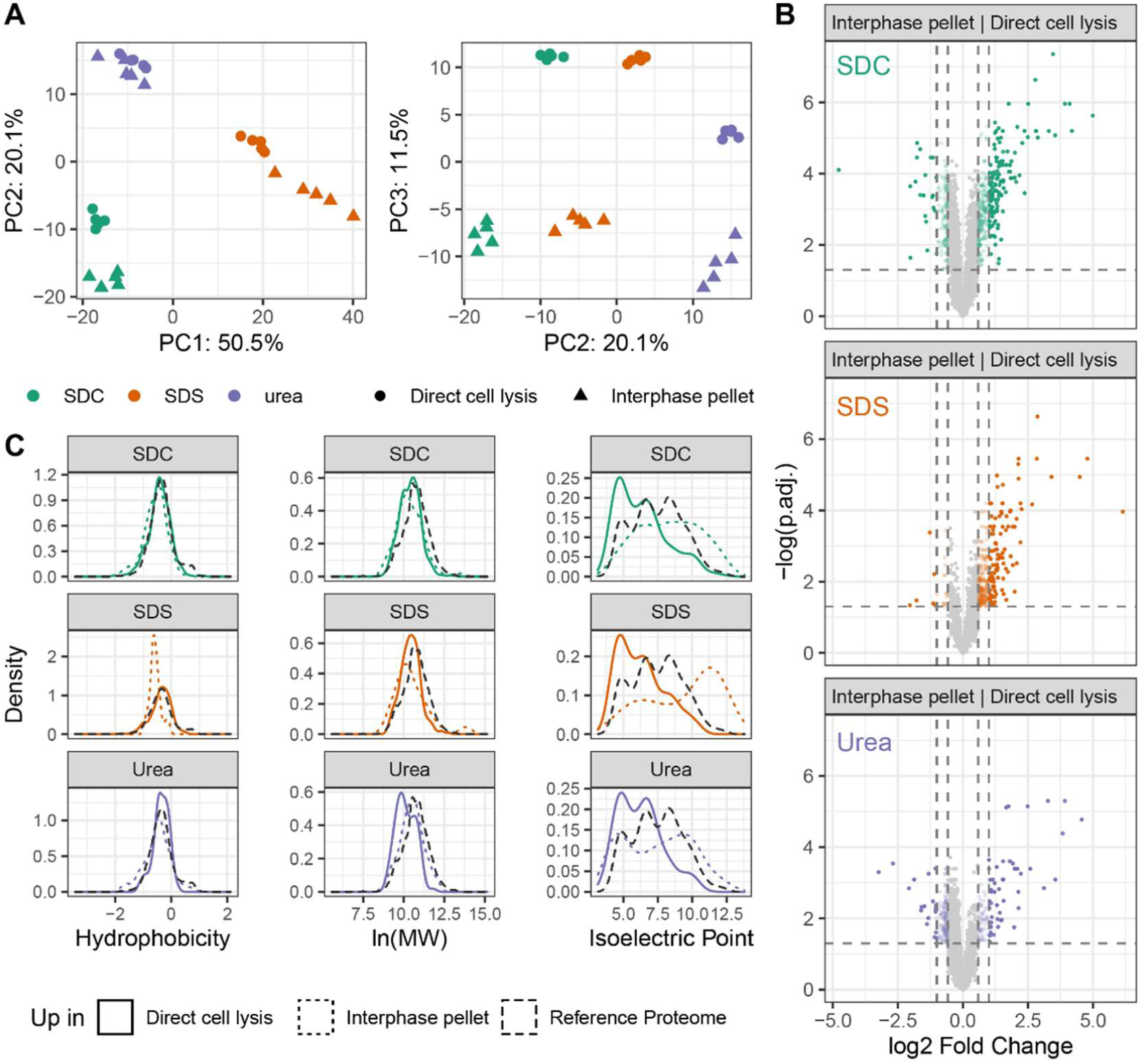
Quantitative comparison of proteins extracted from simultaneous proteo-metabolomics liquid-liquid extraction (SPM-LLE) interphase pellets versus direct cell lysis. A) Principal component analysis of proteins extracted from SPM-LLE interphase pellet (triangles) or direct cell lysis (circles). Protein abundance levels, reproducibly quantified in all independent experiments and conditions, were used as input for PCA. B) Comparison of efficiency to extract proteomes from the SPM-LLE interphase pellets versus direct cell lysis. Significance threshold for enrichment: adjusted *p*-value ≤ 0.05 (two-tailed unpaired *t*-test, Benjamini-Hochberg correction), fold change (FC) of 1.5: colored transparent dots, FC of ≥ 2: colored dots). C) Physicochemical properties of enriched proteins (FC ≥ 1.5, *p*-value ≤ 0.05). Proteins from SPM-LLE interphase pellets are represented with colored dashed lines, proteins from direct cell lysis are represented as colored solid lines. Physicochemical properties of the human reference proteome (SwissProt, uniprot.org) are shown with a black dashed line. Sodium deoxycholate (SDC, green), sodium dodecyl sulfate (SDS, orange), or urea (purple) containing buffer. n= 5 independent experiments (biological replicates).

Extraction by direct cell lysis versus interphase pellets did not elicit differences in hydrophobicity or molecular weight, but the distribution of isoelectric points differed for all extraction agents (Figure 3C). Extraction by direct cell lysis resulted in an enrichment of proteins with lower isoelectric points, whereas proteins with higher isoelectric points were extracted more efficiently from SPM-LLE interphases. This shift is likely explained by the higher coverage of ribosomal and ribonuclear proteins in SPM-LLE interphase pellets (Figure S13, Figure S14). In contrast, proteome extraction by direct cell lysis led to higher coverage of proteins assigned to extracellular exosomes and vesicles (Figure S13).

While a more efficient extraction of proteins that are involved in metabolic pathways was observed with SDS in direct proteome extraction from cells (Figure S15), we did not observe significant differences between SPM-LLE and direct proteome extraction when SDC was used (Figure S16). Urea extracted proteins related to carbohydrate, lipid, and amino acid metabolism more efficiently from the SPM-LLE interphase pellet (Figure S17).

In conclusion, proteome extraction by direct cell lysis yielded higher overall extraction efficiencies for the used surfactants SDC and SDS. In contrast, urea-based solubilisation resulted in higher extraction efficiency of SPM-LLE interphase pellets. All extraction agents solubilized proteins with low isoelectric points more efficiently by direct cell lysis. In contrast, proteins with higher isoelectric points were extracted more efficiently from the SPM-LLE interphase pellets.

## Conclusion

To date, no studies have examined the performance of chaotropes and surfactants for proteome extraction from SPM-LLE interphase pellets and the extent to which proteome coverage and extraction efficiency differ between workup from an SPM-LLE interphase pellet versus direct cell lysis. The studies of Glatter et al. ^12^ and León et al. ^11^ examined the performance of chaotropic agents and surfactants including SDS, SDC and urea for proteome extraction by direct cell lysis from HEK cells and isolated mitochondria. León et al. ^11^ showed that SDS-based extraction was more efficient for protein identification compared to urea for direct cell lysis. This was not the case in our study as SDS overall showed the lowest extraction efficiency for quantitative proteomics for both direct cell lysis and SPM-LLE interphase pellets. One possible explanation for this difference could be different procedures for SDS removal. León et al. used a spin filter-assisted approach to remove SDS, whereas we used the single-pot solid-phase-enhanced sample preparation (SP3) procedure ^13^. The lower extraction efficiency we observed could be assigned to protein losses due to absorption by the carboxylate-modified hydrophilic and hydrophobic beads used for SP3. It should be noted though that despite the overall low performance, SDS-based proteome extraction in combination with SP3 enriched ribosomal proteins and might, therefore, be considered when analysing this specific subproteome. Glatter et al. ^12^ and León et al. ^11^ each concluded that SDC-based extraction was more efficient compared to urea for direct cell lysis. They also showed that tryptic digestion in diluted urea is to be avoided due to high levels of missed cleavages. Although we confirmed this result by comparing the performance of SDC and urea for proteome extraction by direct cell lysis (Figure S17, Table S7), we demonstrated that urea is superior over surfactants for proteome extraction from SPM-LLE interphase pellets. The number of extracted proteins was 3-fold higher for urea as compared to SDC, and 55-fold higher as compared to SDS, with a particularly good performance for the extraction of proteins linked to metabolic pathways. We conclude that of the extraction agents tested here, urea is the most efficient for simultaneous proteo-metabolome analysis.

## ACKNOWLEDGMENT

M.K. thanks the University of Innsbruck (project no: 316826) and the Tyrolian Research Fund (project no: 18903) for financial support. K.T. acknowledges support from the MESI-STRAT project (European Union Horizon 2020 Research and Innovation Program, grant agreement no: 754688), the German Tuberous Sclerosis Foundation, the Stichting TSC Fonds, the PoLiMeR Innovative Training Network (Marie Skłodowska-Curie grant agreement 812616), ARDRE (funded by the EC H2020 Marie Sklodowska-Curie research grant no: 847681), PARC (funded by the European Union’s Horizon Europe research and innovation programme under Grant Agreement no: 101057014, and VERSA (funded by the European’s Union Horizon 2020 Science with & for Society Call (SwafS-2020 Topic 8) under grant agreement no: 101006420).

## Author Contributions

The manuscript was written through contributions of all authors. All authors have given approval to the final version of the manuscript.

## Notes

### Competing Interest Statement

The authors have declared no competing interest.

