## Supporting Information for "Proteome coverage after simultaneous proteo-metabolome liquid-liquid extraction"

| | |
|---|---|
| Supplemental Figures | P. 2-19 |
| Supplemental Tables | P. 2-22 |
| Supplemental Experimental Section | P. 23-27 |
| Supplemental References | P. 27 |

### Supplemental Figures

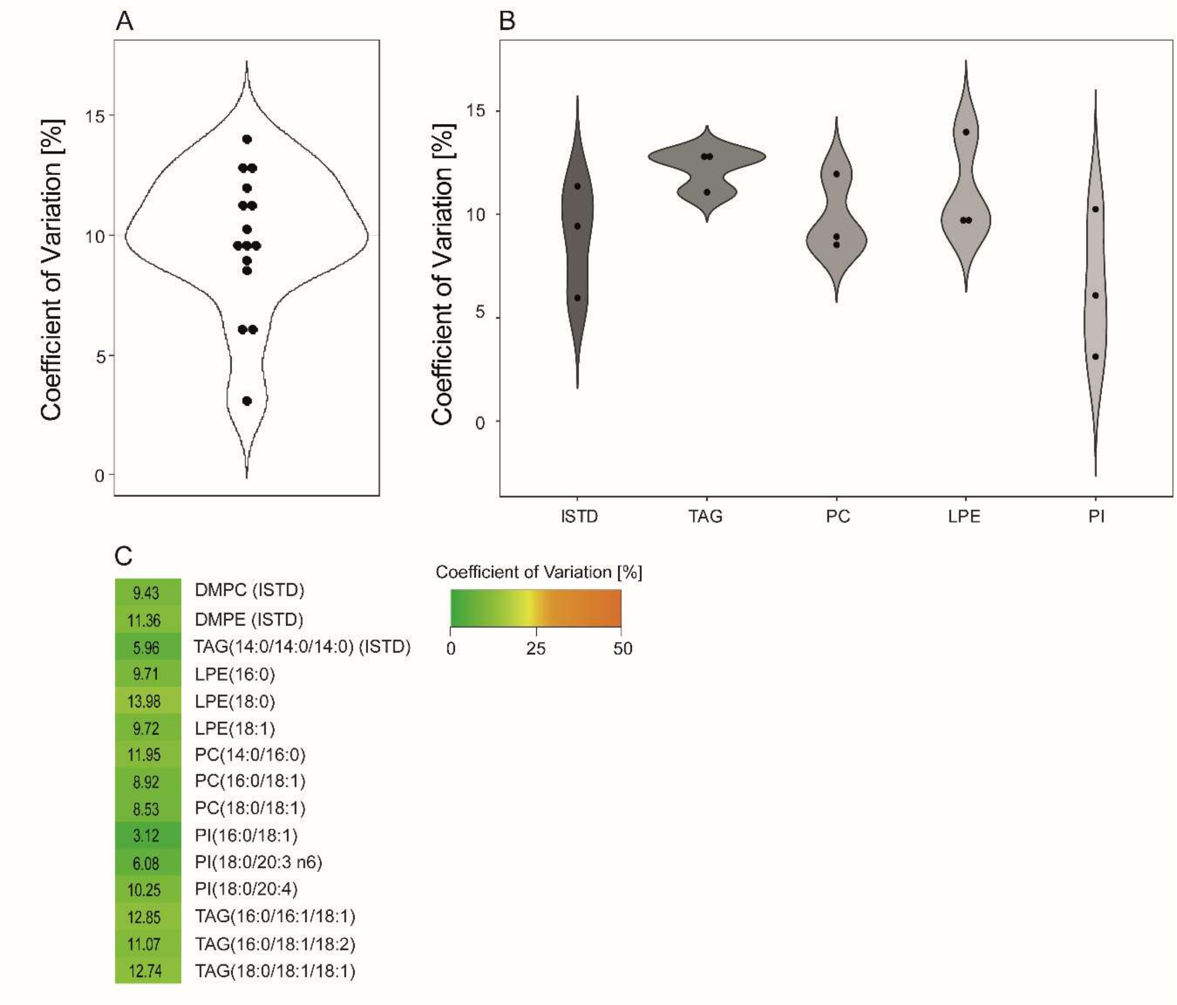


**Figure S1: Quantification of selected lipids extracted from the chloroform phase of a simultaneous proteo-metabolomic liquid-liquid extraction.** Violin plots showing the coefficient of variation [%] for all lipid analytes (A) and sorted based on lipid class and for the internal standards (ISTD, DMPC: 1,2-dimyristoyl-*sn*-glycero-3-phosphocholine, DMPE: 1,2-dimyristoyl-*sn*-glycero-3-phosphorylethanolamine, TMG: 1,2,3-tri-myristoyl-glycerol) (B). C: Colour-coded representation of the coefficient of variation [%] of the different lipid analytes. LPE: lysophosphatidylethanolamine, PC: phosphatidylcholine, PI: phosphatidylinositol, TAG: triglyceride. n= 5 independent experiments (biological replicates).

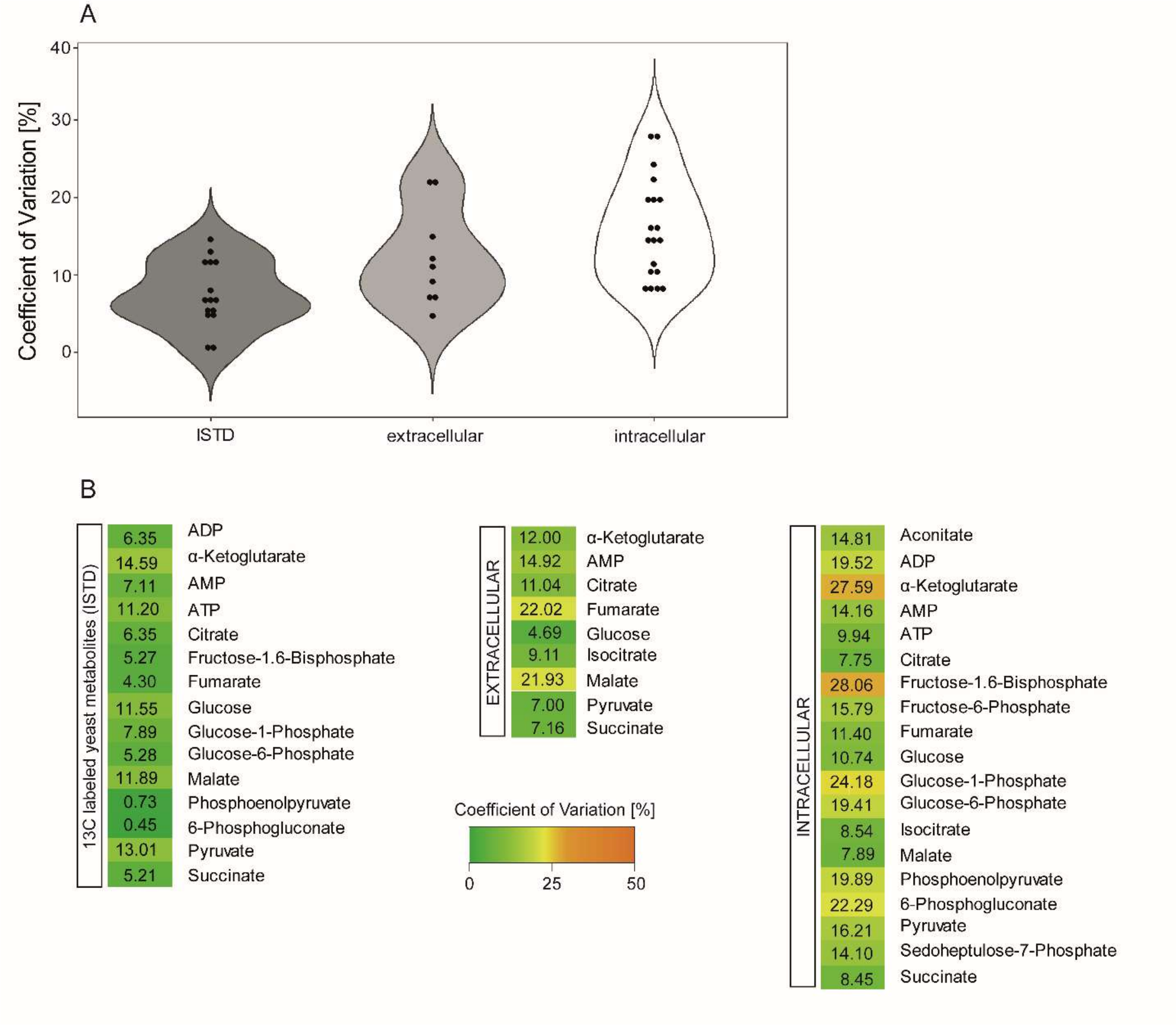


**Figure S2: Quantification of polar metabolites extracted from the methanol phase of a simultaneous proteo-metabolomics liquid-liquid extraction.** A: Violin plots showing the coefficient of variation [%] for internal standards (ISTD), extracellular and intracellular polar metabolites. B: Colour-coded representation of the coefficient of variation [%] of the $^{13}C$ labelled yeast metabolites used as internal standards (ISTD), extracellular and intracellular polar metabolites. n= 5 independent experiments (biological replicates).

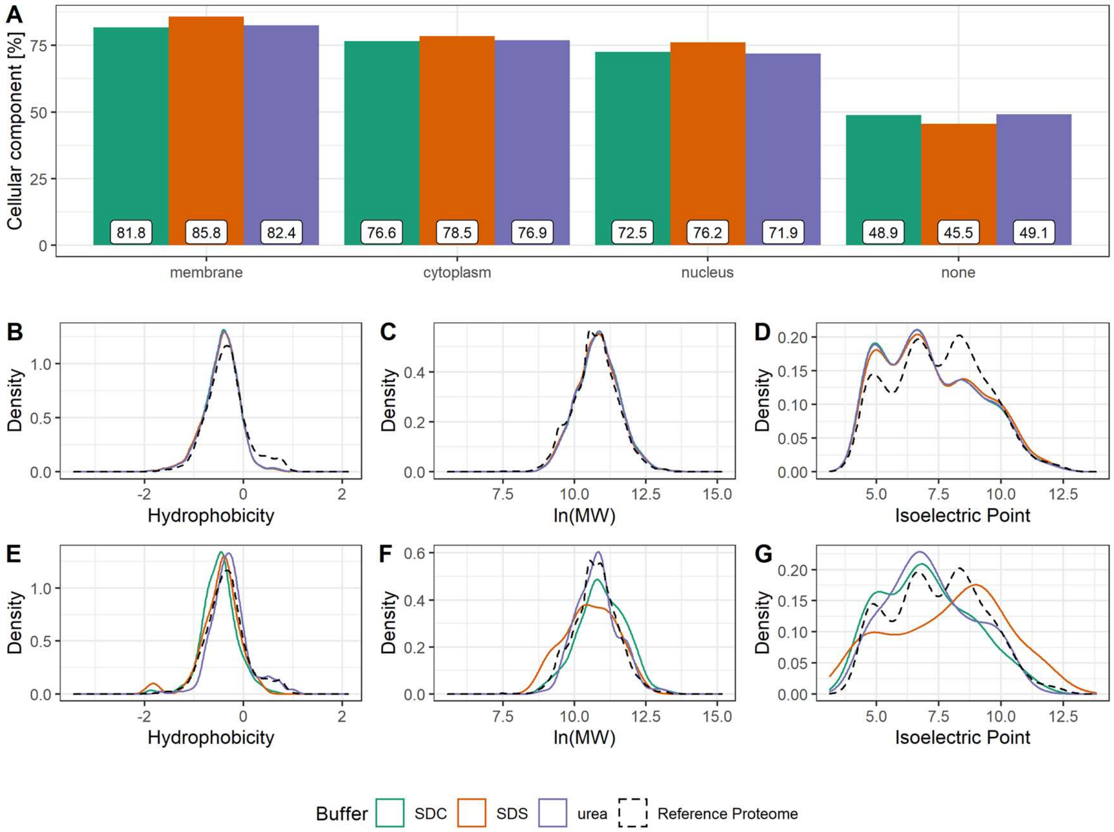


**Figure S3: Localization and physicochemical properties of proteins identified in simultaneous proteo-metabolomic liquid-liquid extraction (SPM-LLE) interphase pellets.** A: Distribution of proteins across the main GO cellular component categories (membrane proteins, GO: 0016020; nuclear proteins, GO: 0005634; cytoplasmatic proteins, GO: 0005737). B, C, D: Physicochemical properties (hydrophobicity (B), molecular weight (C), isoelectric point (D)) of all proteins identified with the three extractants (sodium deoxycholate (SDC, green), sodium dodecyl sulfate (SDS, orange), urea (purple)). E, F, G: Physicochemical properties (hydrophobicity (E), molecular weight (F), isoelectric point (G)) of proteins identified exclusively upon extraction with SDS (orange), SDC (green) or urea (blue) extracts. Dashed line, human reference proteome (SwissProt, uniprot.org).

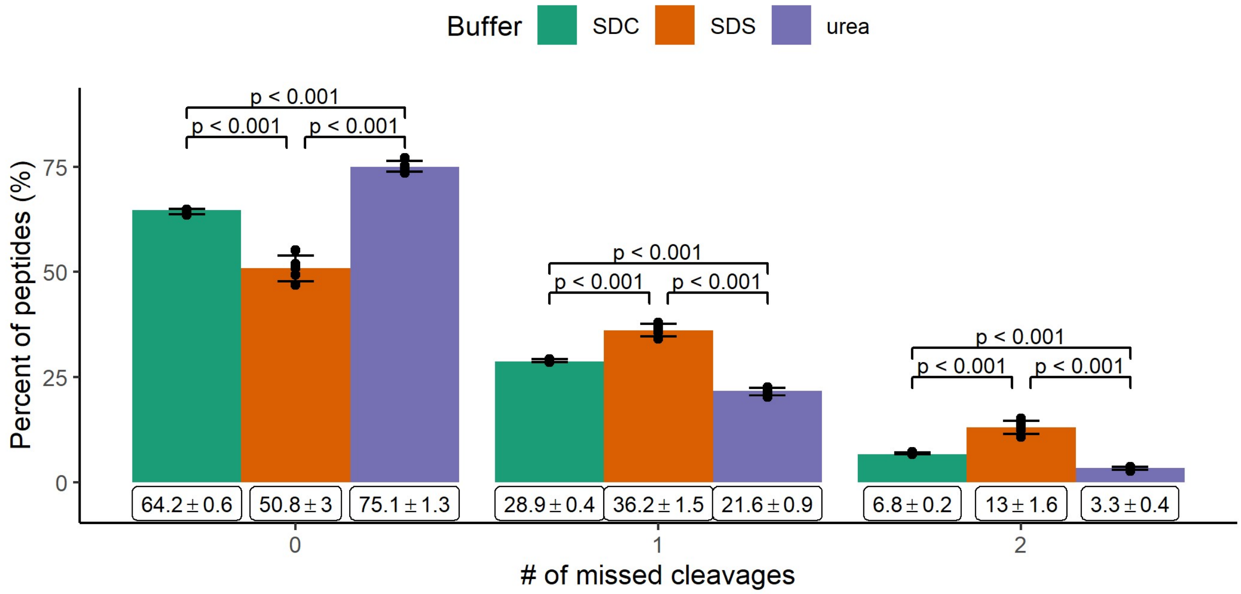


**Figure S4: Relative percentage of missed cleavages after tryptic digestion of the interphase pellet extracted by sodium deoxycholate (SDC, green), sodium dodecyl sulfate (SDS, orange), urea (purple) based buffer systems.** Bar graph: mean with standard deviation, Statistical analysis: two-tailed unpaired *t*-test. n= 5 independent experiments (biological replicates).

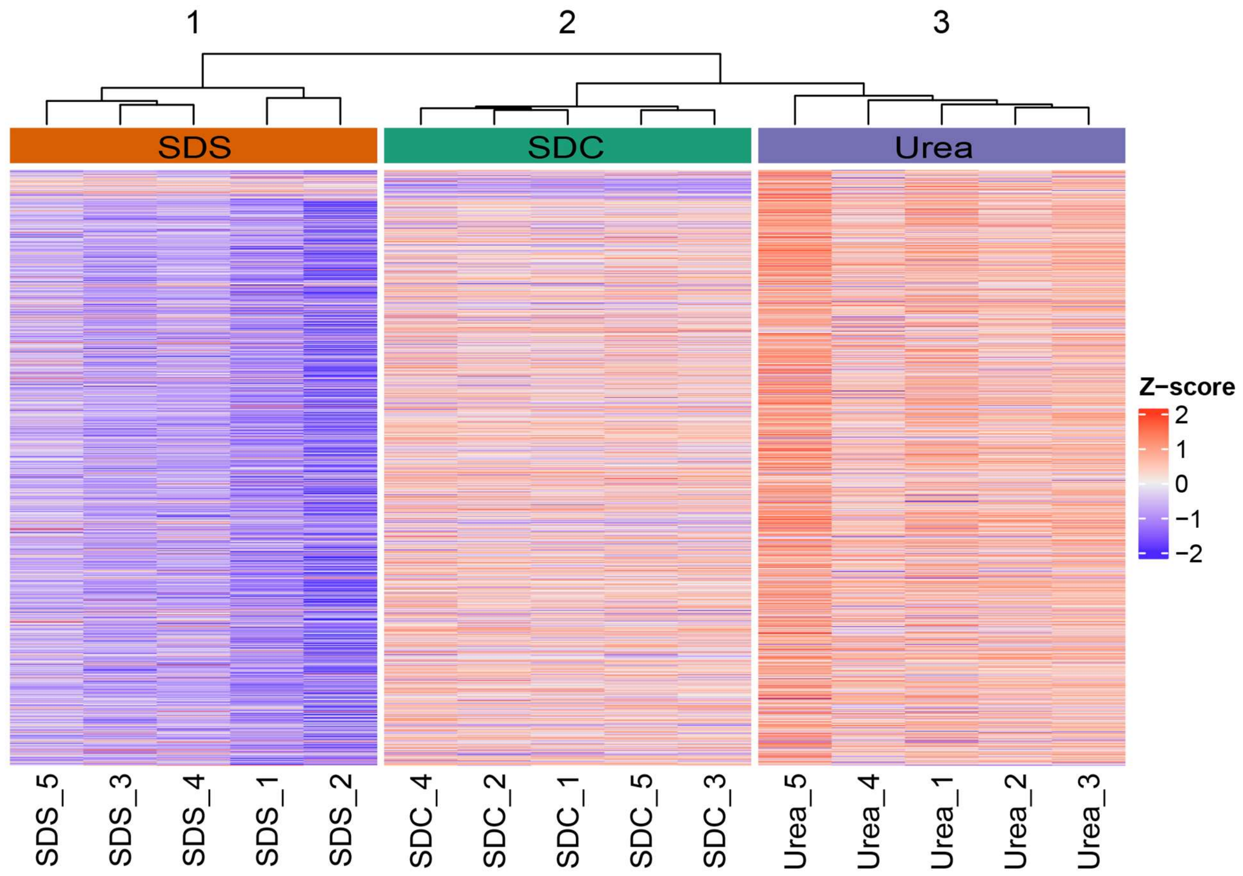


**Figure S5: Relative abundances of all proteins quantified in simultaneous proteo-metabolomics liquid-liquid extraction interphase pellets extracted by sodium deoxycholate (SDC, green), sodium dodecyl sulfate (SDS, orange), urea (purple) based buffer systems.** n= 5 independent experiments (biological replicates).

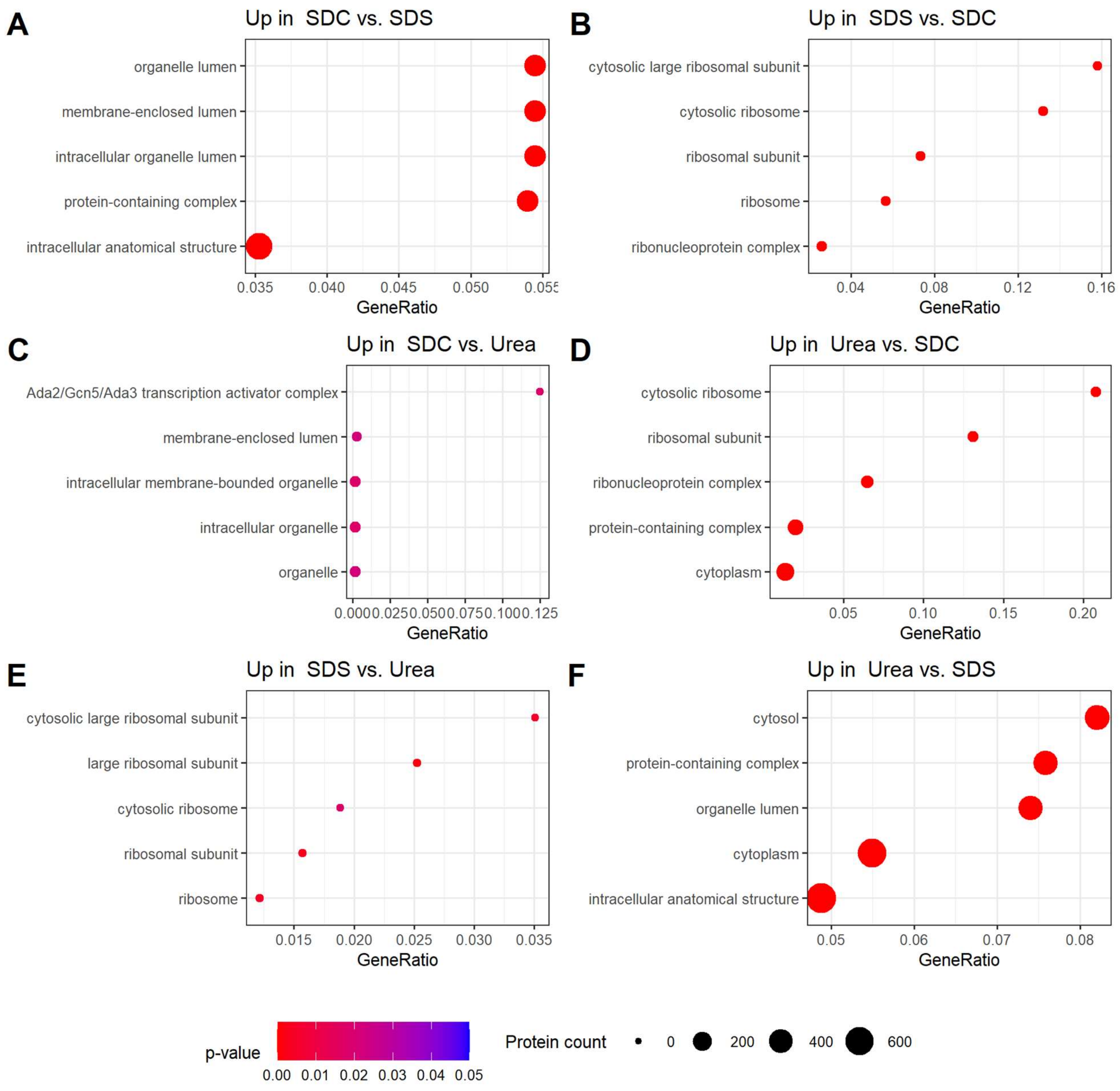


**Figure S6: Gene-Ontology (GO)-enrichment (cellular component) of proteins extracted more efficiently with sodium deoxycholate (SDC), sodium dodecyl sulfate (SDS) or urea.** Proteins are considered to be extracted more efficiently if they showed a fold-change of ≥ 1.5 and an adjusted *p*-value ≤ 0.05 (see Figure 3B). The size of the dots indicates the protein count of enriched proteins related to the specific GO-term. n= 5 independent experiments (biological replicates).

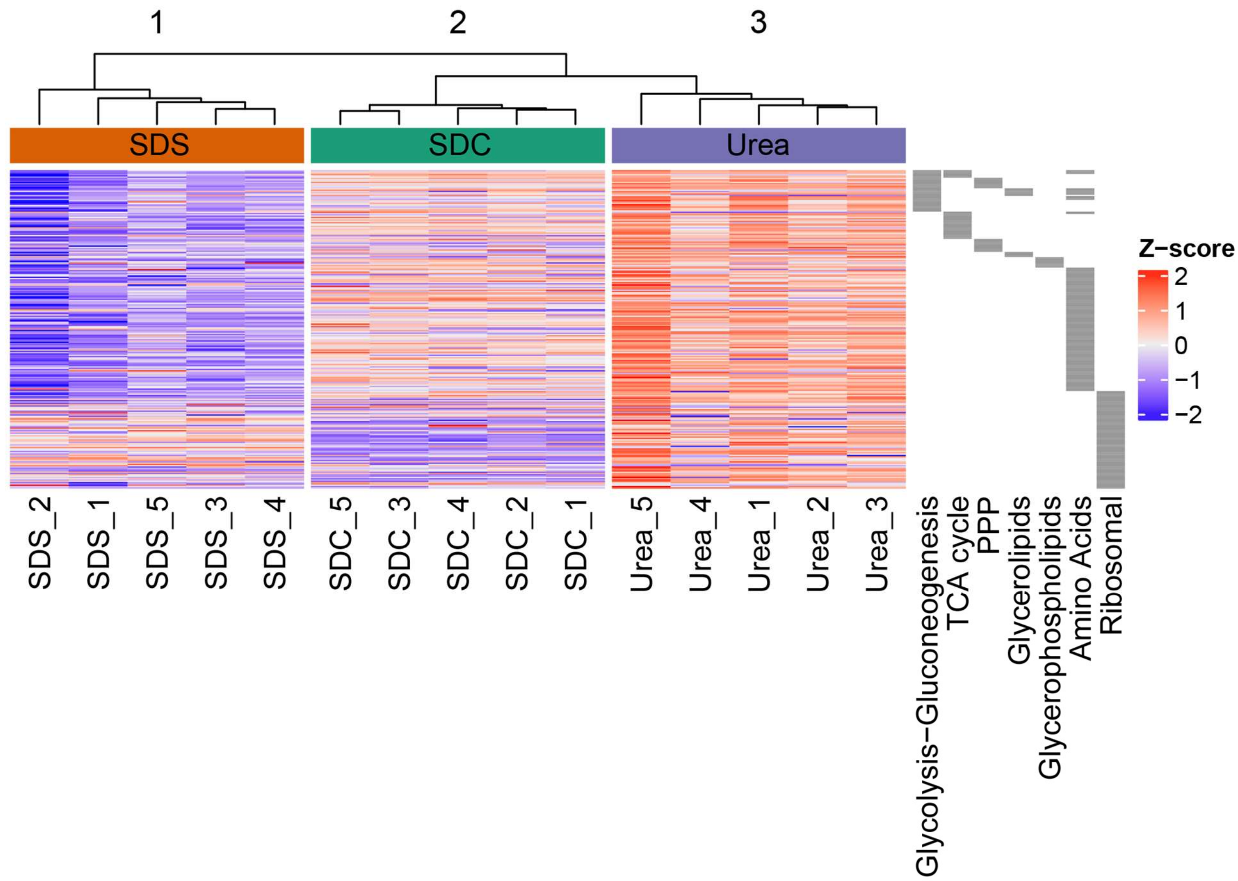


**Figure S7: Relative abundancies related to specific KEGG pathways for proteins identified from interphase pellets extracted with SDS (green), SDC (orange) or urea (purple).** Glycolysis-Gluconeogenesis: hsa00010; TCA cycle: hsa00020; Phosphate Pentose Pathway (PPP): hsa00030; Glycerolipids: hsa00561; Glycerophospholipids: hsa00564; Amino Acids: hsa00220, hsa00250, hsa00260, hsa00270, hsa00280, hsa00290, hsa00300, hsa00310, hsa00330, hsa00340, hsa00350, hsa00360, hsa00380, hsa00400; Ribosomal Proteins: hsa03010. n= 5 independent experiments (biological replicates).

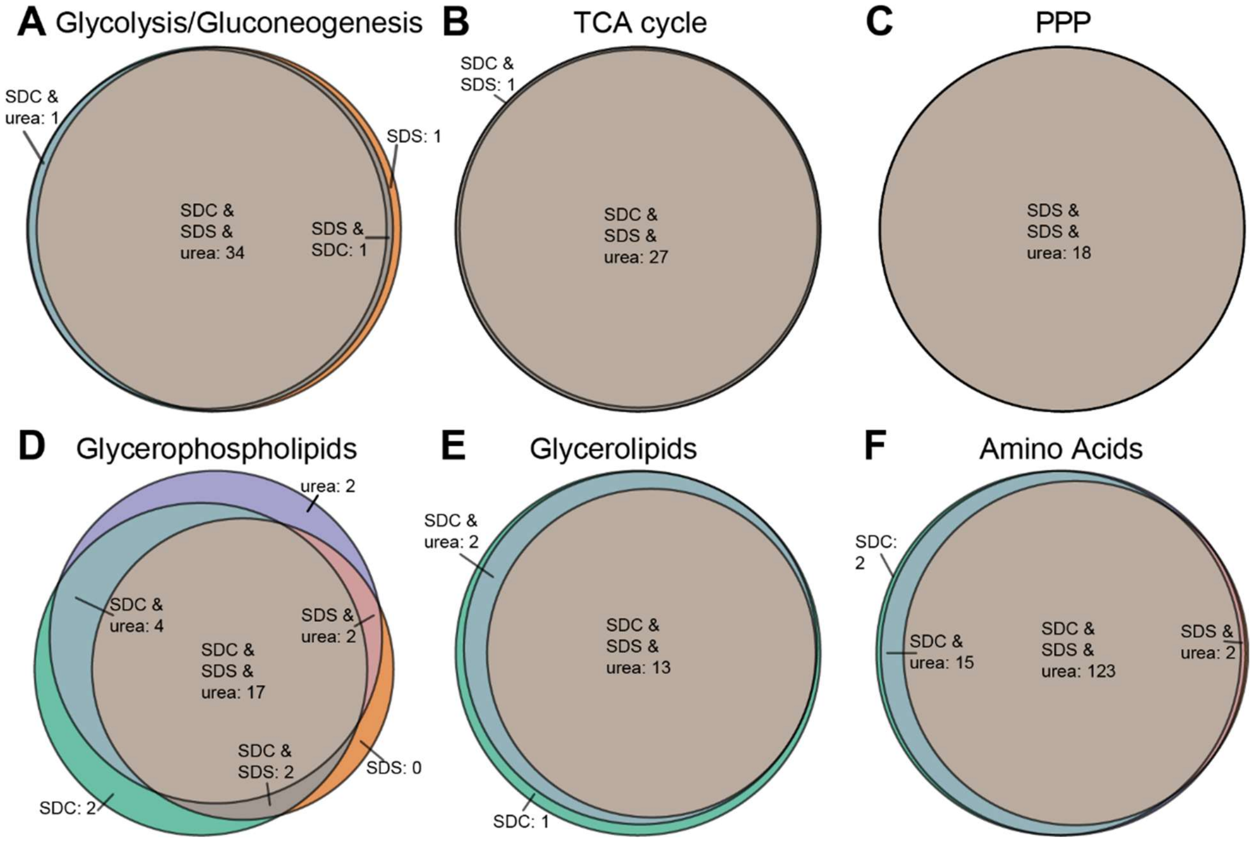


**Figure S8: Comparison of identified proteins that are related to metabolic pathways.** Number of proteins reproducibly identified in all independent experiments (n = 5, biological replicates).

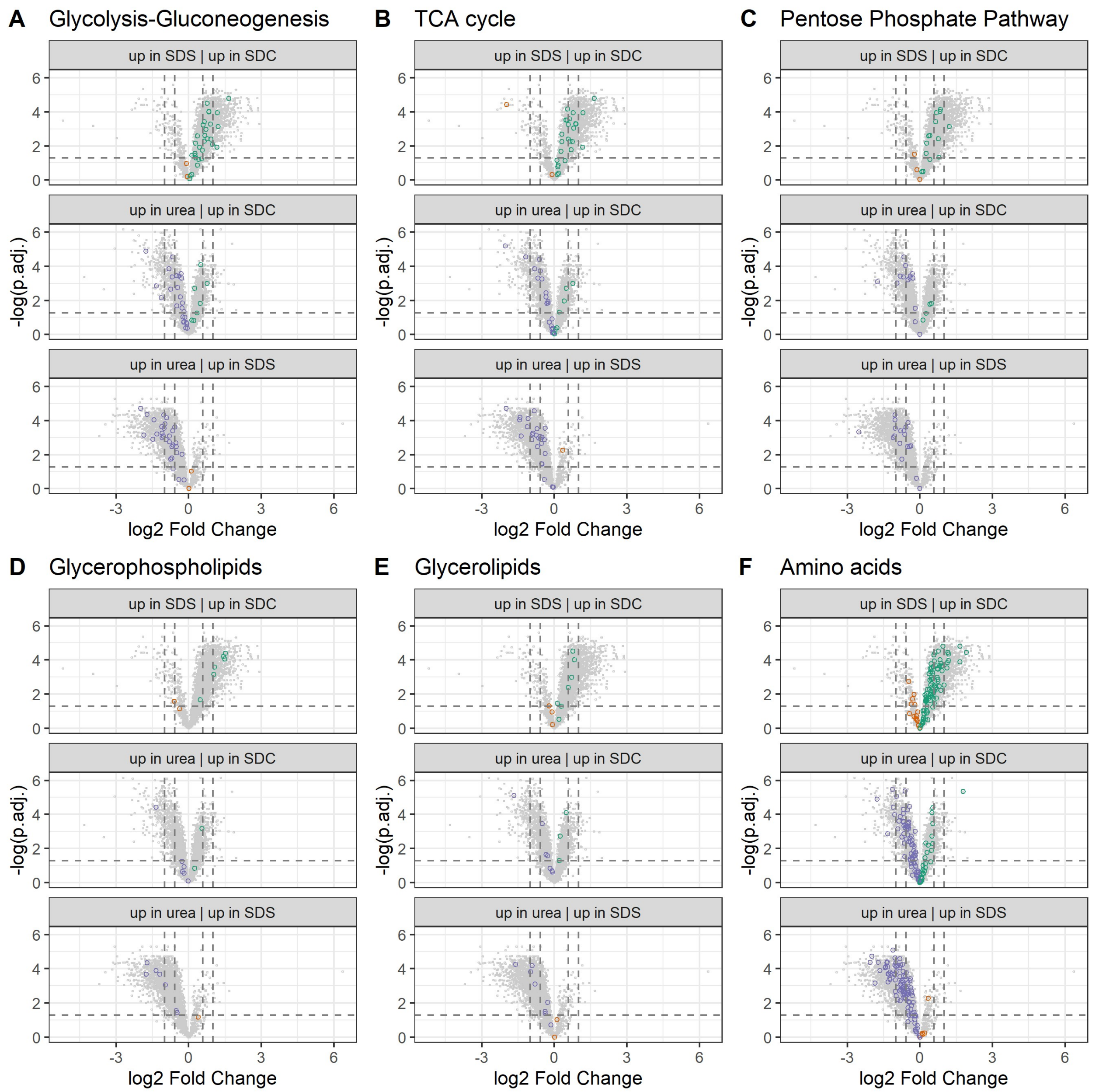


**Figure S9: Extraction efficiencies for proteins related to metabolic pathways (KEGG) from interphase pellets extracted with sodium deoxycholate (SDC), sodium dodecyl sulfate (SDS) or urea**. Proteins that are associated with glycolysis-gluconeogenesis (hsa00010) TCA cycle (hsa00020), pentose phosphate pathway (PPP) (hsa00030), glycerolipids (hsa00561), glycerophospholipids (hsa00564), amino acids (hsa00220, hsa00250, hsa00260, hsa00270, hsa00280, hsa00290, hsa00300, hsa00310, hsa00330, hsa00340, hsa00350, hsa00360, hsa00380, hsa00400) are highlighted dependent on the extraction agent in purple (urea), green (SDC) and orange (SDS). Vertical line: threshold adjusted *p*-value: 0.05. Two horizontal lines: fold change (FC) threshold: 1.5 and >2. n= 5 independent experiments (biological replicates).

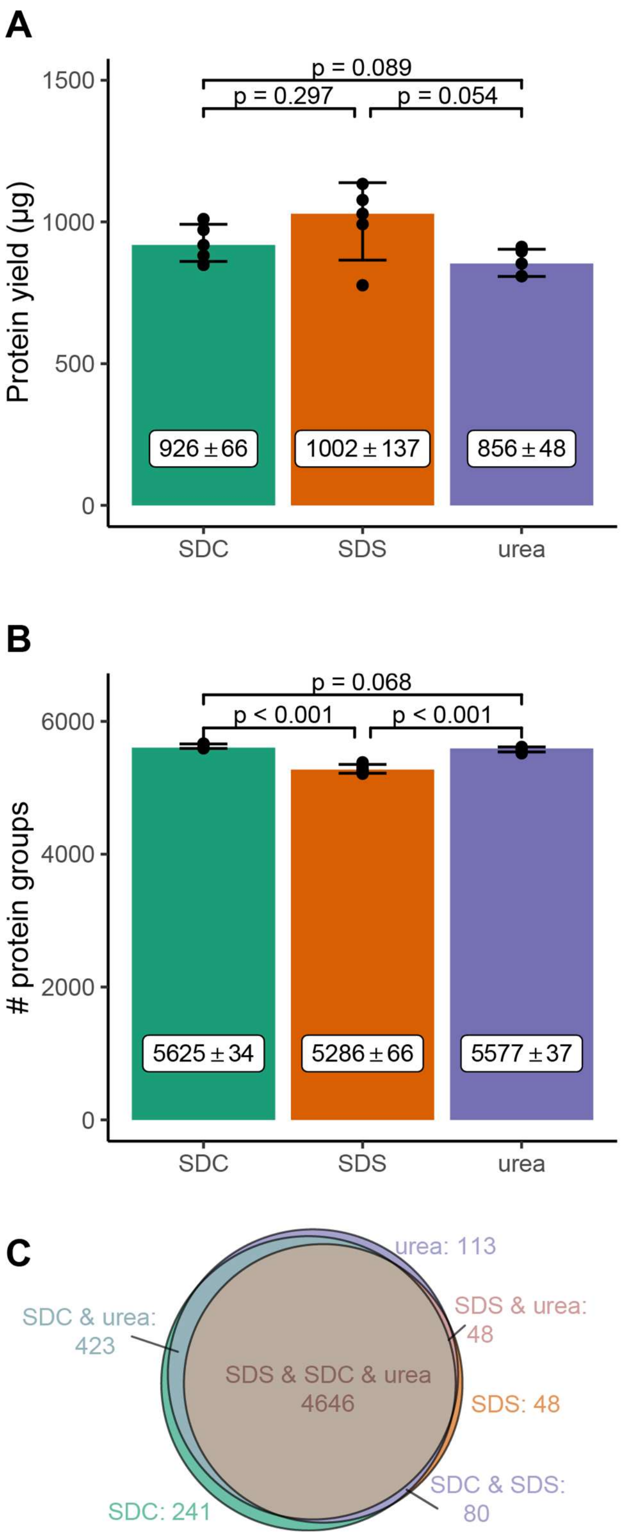


**Figure S10: Protein yield (A) and number of proteins identified (B, C) by direct cell lysis.** A) Protein yield (µg) and B) number of identified proteins by direct cell lysis using sodium deoxycholate (SDC), sodium dodecyl sulfate (SDS), and urea. Bar graph: mean with standard deviation. Statistical analyses with two-tailed unpaired *t*-test. C) Number of proteins reproducibly identified in all biological replicates (n = 5) with the indicated extraction agents. Average number of cells: 4.3 x $10^6$ cells.

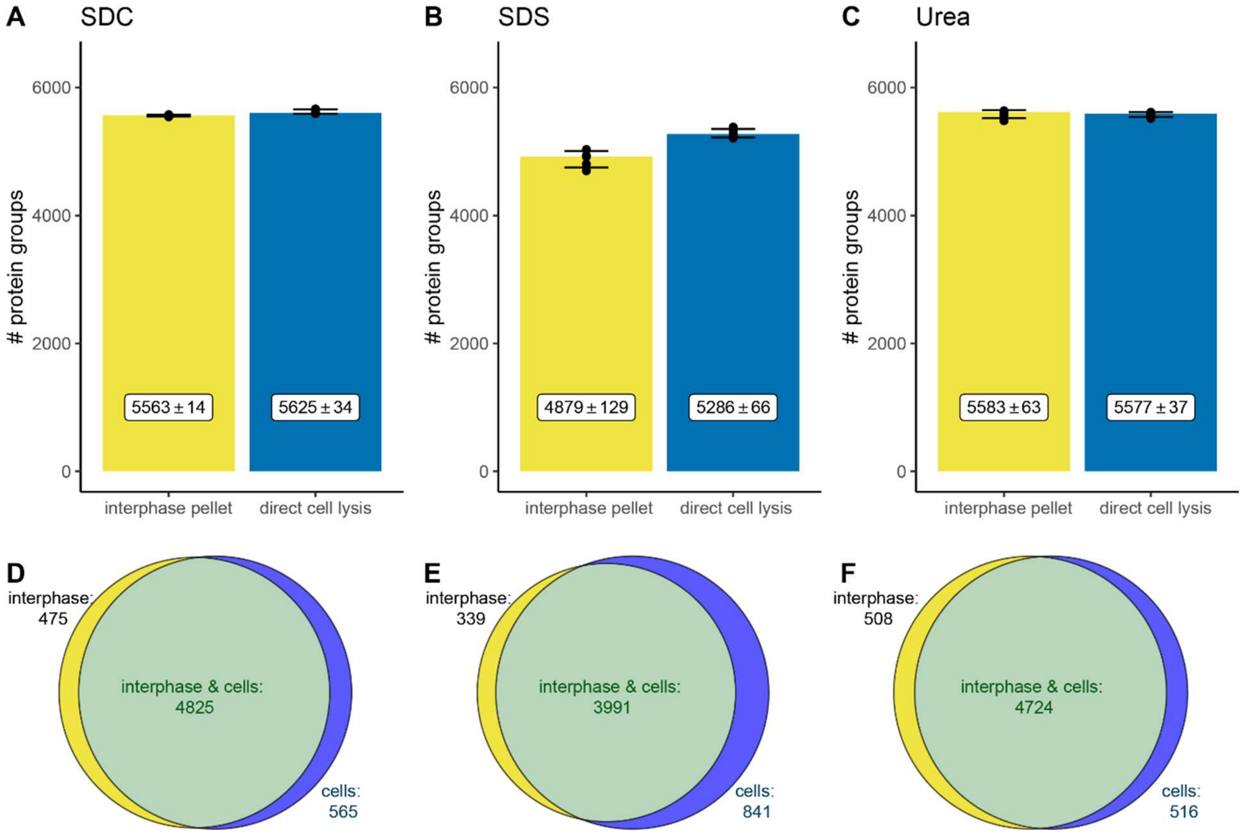


**Figure S11: Proteins identified by proteome extraction from interphases of SPM-LLE and by direct cell lysis.** A-C: Number of identified proteins in the proteomes extracted from interphases of SPM-LLE (same data as shown in Figure 1 B) and by direct cell lysis (same data as shown in Figure S10 B) using SDC (A), SDS (B) and urea (C). Mean with standard deviation. D-F: C: Venn diagrams showing the number of proteins reproducibly identified in all replicates using SDC (D), SDS (E) and urea (F). Statistical analyses were performed using the two-tailed unpaired *t*-test. n= 5 independent experiments (biological replicates).

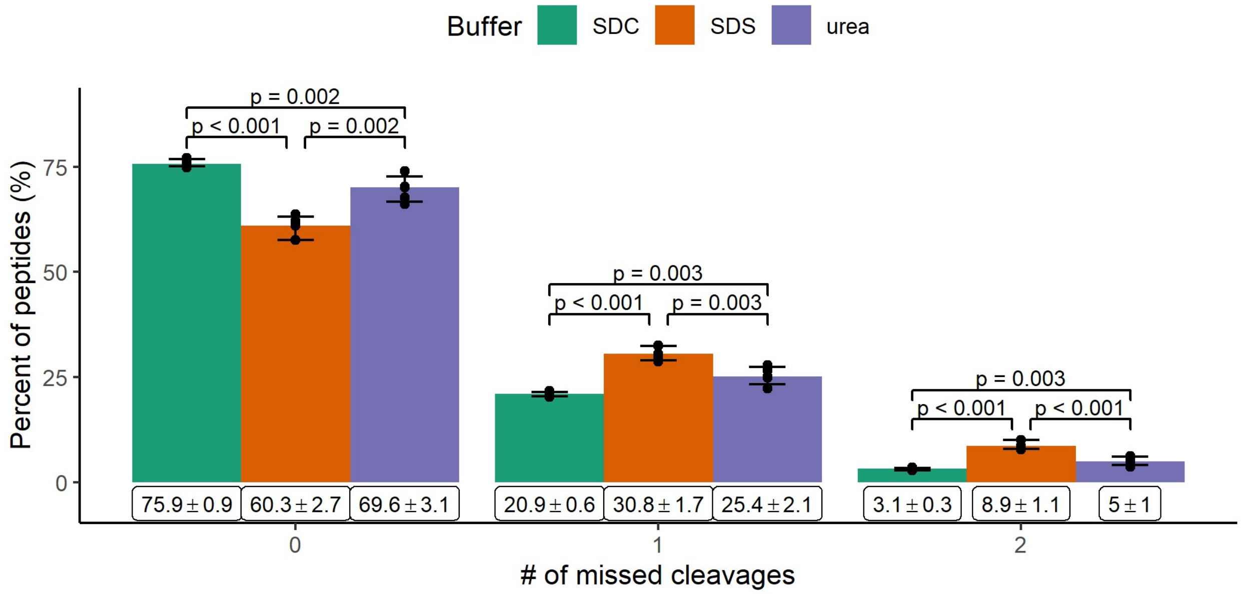


**Figure S12: Analysis of missed cleavages during tryptic digestion of proteomes obtained by direct cell lysis using SDC (green), SDS (orange) and urea (purple).** Relative percentage of missed cleavages during tryptic digestion using SDC, SDS and urea, mean with standard deviation. Statistical analyses were performed using the two-tailed unpaired *t*-test. n= 5 independent experiments (biological replicates).

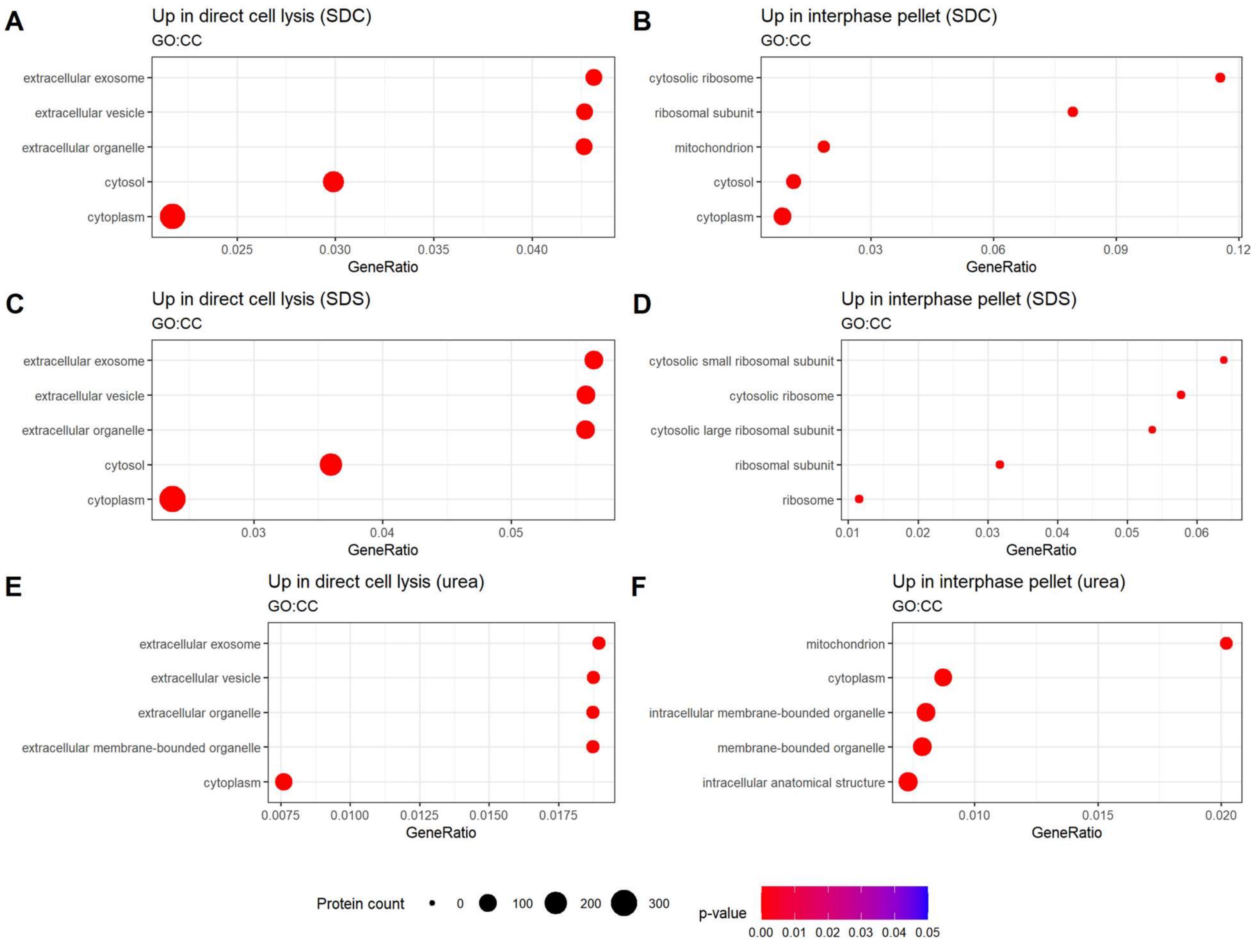


**Figure S13: Gene Ontology (GO) enrichment analysis of cellular component (GO:CC) of proteins extracted significantly more efficiently by direct cell lysis (up in cells) versus SPM-LLE interphase pellet.** Proteins were considered extracted significantly more efficiently enriched with a FC ≥ 2 and a *p*-value ≤ 0.05. The size of the dots indicates the number of enriched proteins related to the specific GO-term. n= 5 independent experiments (biological replicates).

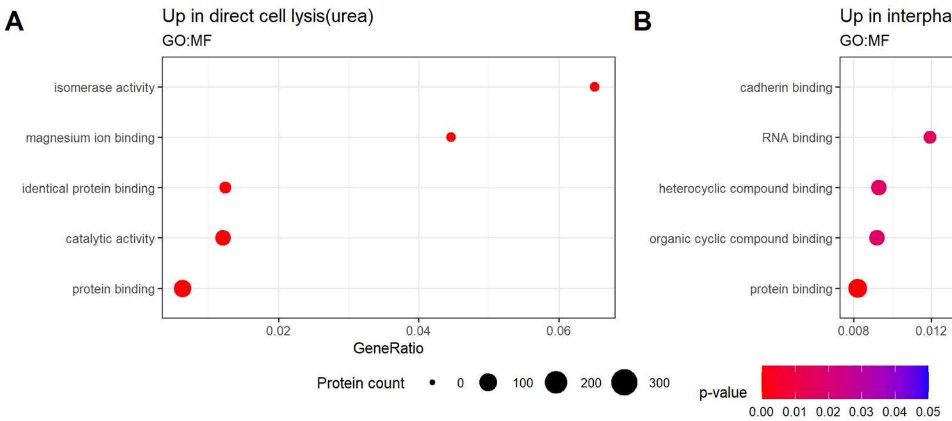


**Figure S14: Gene Ontology (GO) enrichment analysis of molecular function (GO:MF) of proteins extracted significantly more efficiently by direct cell lysis (up in cells) versus SPM-LLE interphase pellet using urea.** Proteins were considered extracted significantly more efficiently enriched with a FC ≥ 2 and a *p*-value ≤ 0.05. The size of the dots indicates the number of enriched proteins related to the specific GO-term. n= 5 independent experiments (biological replicates).

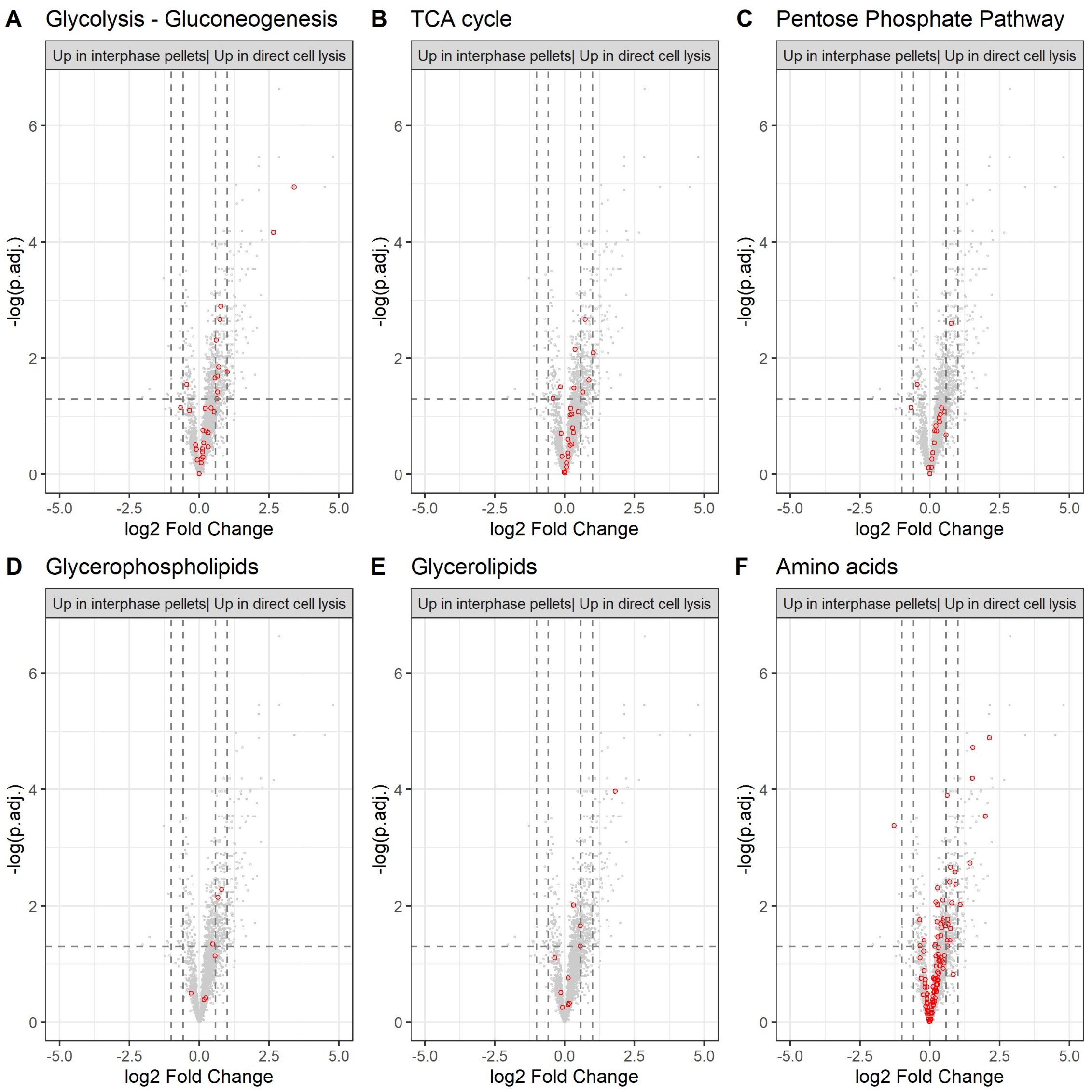


**Figure S15: Differential analysis of extraction efficiencies for protein related to metabolic pathways (KEGG) using SDS to solubilize SPM-LLE interphase pellets (interphases) versus direct cell lysis (cells).** Proteins that are associated with glycolysis-gluconeogenesis (hsa00010) TCA cycle (hsa00020), pentose phosphate pathway (PPP) (hsa00030), glycerolipids (hsa00561), glycerophospholipids (hsa00564), amino acids (hsa00220, hsa00250, hsa00260, hsa00270, hsa00280, hsa00290, hsa00300, hsa00310, hsa00330, hsa00340, hsa00350, hsa00360, hsa00380, hsa00400) are highlighted in red circles. Vertical line: threshold adjusted *p*-value: 0.05. Two horizontal lines: fold change (FC) threshold: 1.5 and >2. n= 5 independent experiments (biological replicates).

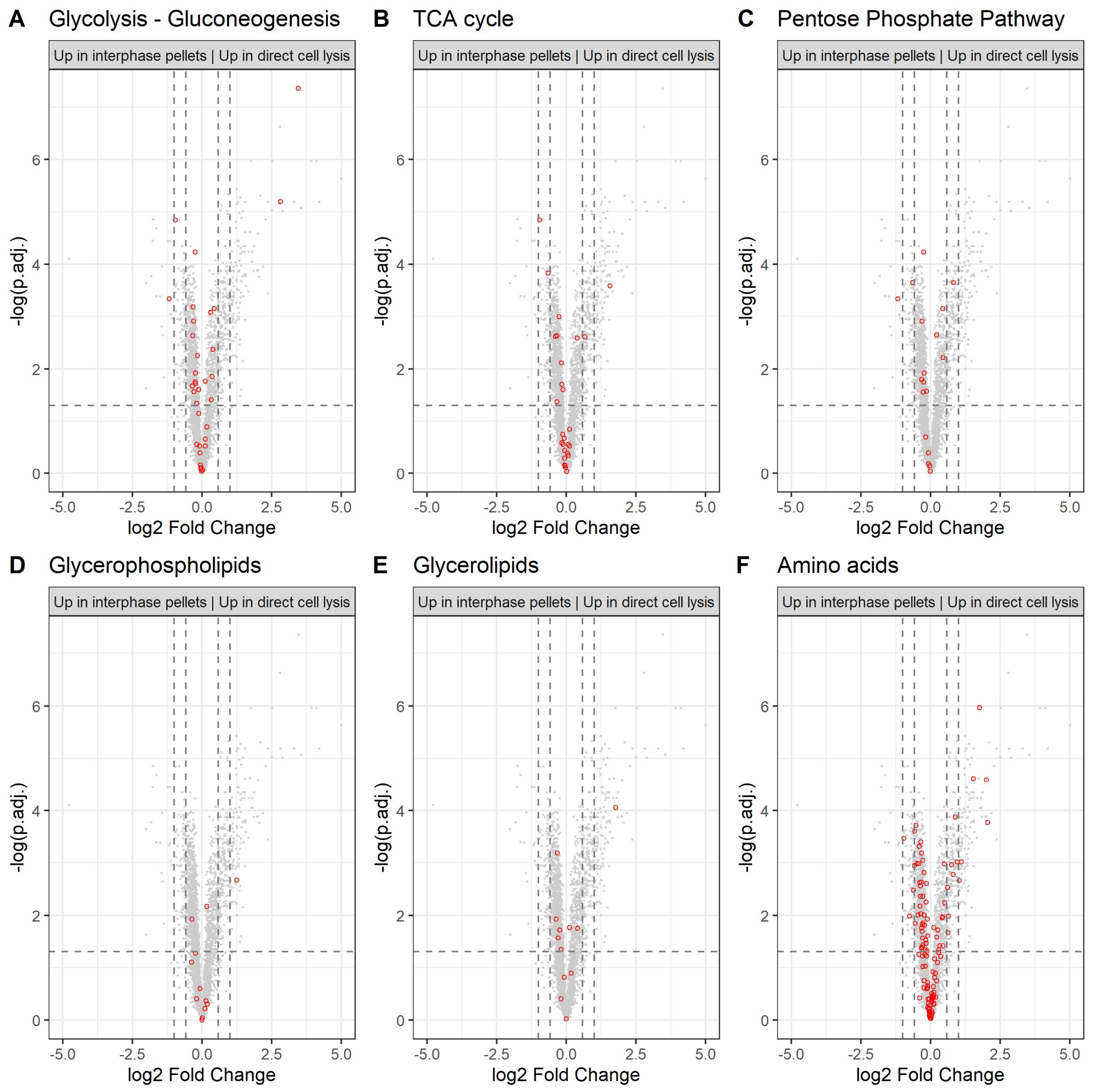


**Figure S16: Differential analysis of extraction efficiencies for proteins related to metabolic pathways using SDC to solubilize SPM-LLE interphase pellets (interphases) versus direct cell lysis (cells).** Proteins that are associated with glycolysis-gluconeogenesis (hsa00010) TCA cycle (hsa00020), pentose phosphate pathway (PPP) (hsa00030), glycerolipids (hsa00561), glycerophospholipids (hsa00564), amino acids (hsa00220, hsa00250, hsa00260, hsa00270, hsa00280, hsa00290, hsa00300, hsa00310, hsa00330, hsa00340, hsa00350, hsa00360, hsa00380, hsa00400) are highlighted in red circles. Vertical line: threshold adjusted *p*-value: 0.05. Two horizontal lines: fold change (FC) threshold: 1.5 and >2. n= 5 independent experiments (biological replicates).

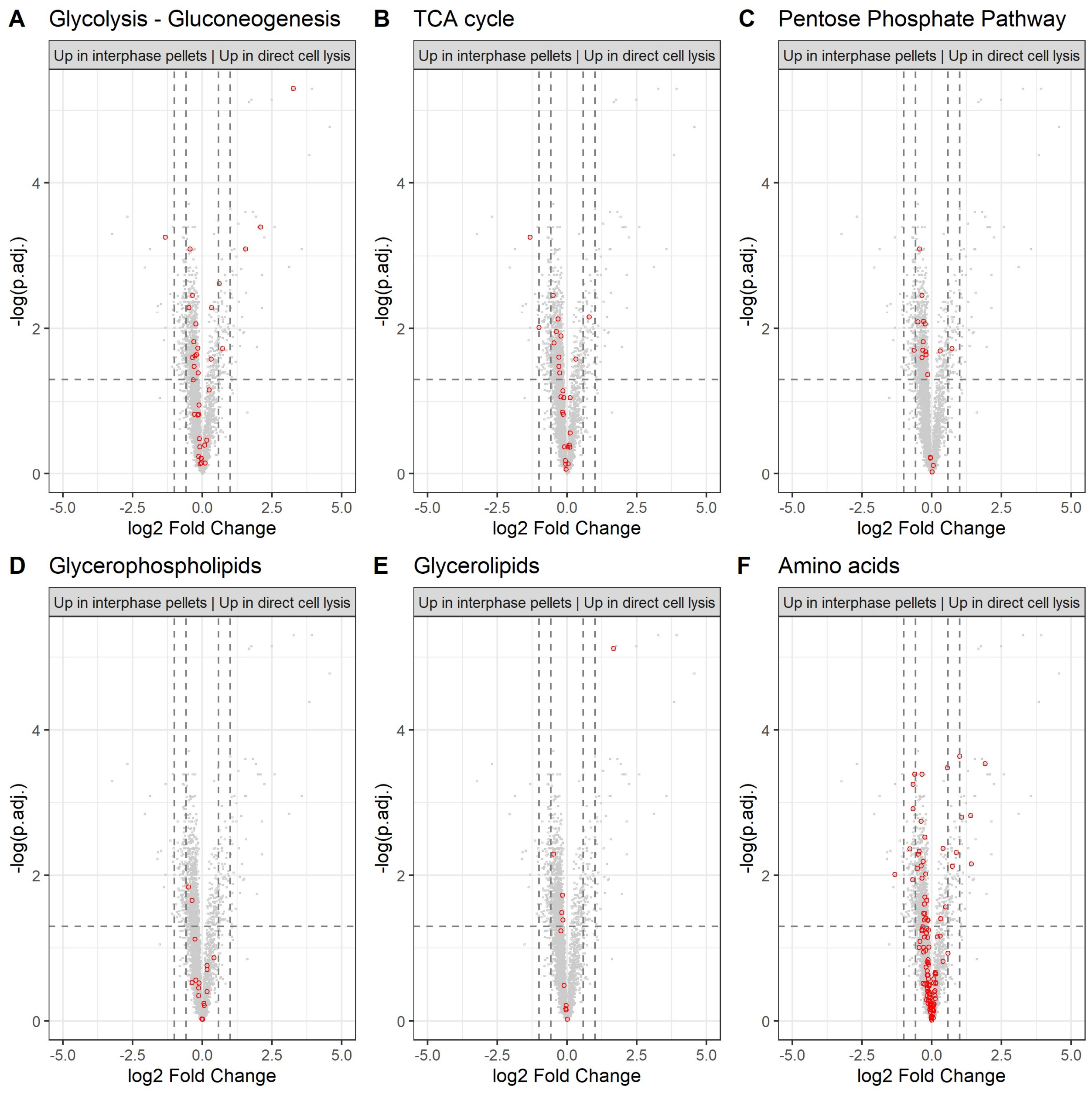


**Figure S17: Differential analysis of extraction efficiencies for proteins related to metabolic pathways using urea to solubilize SPM-LLE interphase pellets (interphases) versus direct cell lysis (cells).** Proteins that are associated with glycolysis-gluconeogenesis (hsa00010) TCA cycle (hsa00020), pentose phosphate pathway (PPP) (hsa00030), glycerolipids (hsa00561), glycerophospholipids (hsa00564), amino acids (hsa00220, hsa00250, hsa00260, hsa00270, hsa00280, hsa00290, hsa00300, hsa00310, hsa00330, hsa00340, hsa00350, hsa00360, hsa00380, hsa00400) are highlighted in red circles. Vertical line: threshold adjusted *p*-value: 0.05. Two horizontal lines: fold change (FC) threshold: 1.5 and >2. n= 5 independent experiments (biological replicates).

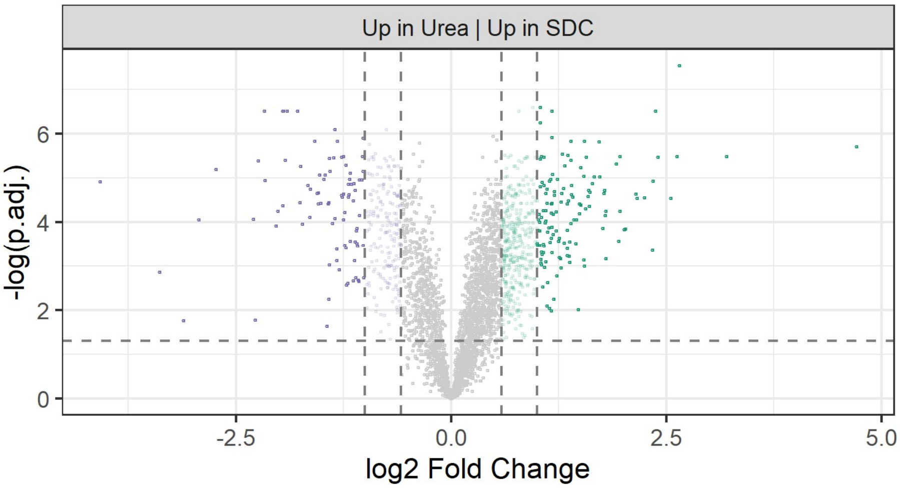


**Figure S18: Quantitative comparison of proteins extracted by direct cell lysis using urea (purple) or SDC (green).** Significance threshold for enrichment: adjusted *p*-value ≤ 0.05 (two-tailed unpaired *t*-test, Benjamini-Hochberg correction), fold change (FC) of 1.5: colored transparent dots, FC of ≥ 2: colored dots).

### Supplemental Tables

**Table S1**: Number of reproducibly identified proteins in the proteomes extracted by urea, sodium deoxycholate (SDC) and sodium dodecyl sulfate (SDS) from the SPM-LLE interphase protein pellet. Urea+SDC+SDS: number of identified proteins identified in both urea, SDC and SDS. Urea+SDC: number of identified proteins identified in both urea and SDC. Urea+SDS: number of identified proteins identified in both urea and SDS. SDC+SDS: number of identified proteins identified in both SDC and SDS. Urea: number of proteins identified in urea only. SDC: number of proteins identified in SDC only. SDS: number of proteins identified in SDS only.

| Condition | Number of identified protein groups | Percentage |
|---|---|---|
| Urea+SDC+SDS | 4200 | 75.5 |
| Urea+SDC | 795 | 14.3 |
| Urea+SDS | 38 | 0.7 |
| SDC+SDS | 63 | 1.1 |
| Urea | 199 | 3.6 |
| SDC | 242 | 4.3 |
| SDS | 29 | 0.5 |

**Table S2**: Relative percentage of missed cleavages during tryptic digestion after proteome extraction from SPM-LLE interphase pellet (SPM-LLE interphase pellet) and after proteome extraction by direct cell lysis (Direct cell lysis) using sodium deoxycholate (SDC).

| Number of missed cleavages | SPM-LLE interphase pellet [%] | Direct cell lysis [%] |
|---|---|---|
| 0 | 64.2 ±0.6 | 75.9 ±0.9 |
| 1 | 28.9 ±0.4 | 20.9 ±0.6 |
| 2 | 6.8 ±0.2 | 3.1 ±0.3 |

**Table S3**: Relative percentage of missed cleavages during tryptic digestion after proteome extraction from SPM-LLE interphase pellet (SPM-LLE interphase pellet) and after proteome extraction by direct cell lysis (Direct cell lysis) using sodium dodecyl sulfate (SDS).

| Number of missed cleavages | SPM-LLE interphase pellet [%] | Direct cell lysis [%] |
|---|---|---|
| 0 | 50.8 ±3 | 60.3 ±2.7 |
| 1 | 36.2 ±1.5 | 30.8 ±1.7 |
| 2 | 13 ±1.6 | 8.9 ±1.1 |

**Table S4**: Relative percentage of missed cleavages during tryptic digestion after proteome extraction from SPM-LLE interphase pellet (SPM-LLE interphase pellet) and after proteome extraction by direct cell lysis (Direct cell lysis) using urea.

| Number of missed cleavages | SPM-LLE interphase pellet [%] | Direct cell lysis [%] |
|---|---|---|
| 0 | 75.1 ±1.3 | 69.6 ±3.1 |
| 1 | 21.6 ±0.9 | 25.4 ±2.1 |
| 2 | 3.3 ±0.4 | 5 ±1 |

**Table S5:** Number of proteins related to metabolic pathways extracted from SPM-LLE interphase pellet with at least 1.5-fold higher efficiency.

| | SDS vs. SDC | | urea vs. SDC | | urea vs. SDS | |
|---|---|---|---|---|---|---|
| | # proteins up in SDS | # proteins up in SDC | # proteins up in urea | # proteins up in SDC | # proteins up in urea | # proteins up in SDS |
| Glycolysis/gluconeogenesis | 0 | 16 | 7 | 1 | 22 | 0 |
| Tricarboxylic acid cycle | 1 | 11 | 5 | 1 | 15 | 0 |
| Pentose phosphate pathway | 0 | 7 | 6 | 0 | 11 | 0 |
| Glycerophospholipids metabolism | 1 | 5 | 1 | 0 | 5 | 0 |
| Glycerolipid metabolism | 0 | 3 | 1 | 0 | 4 | 0 |
| Amino acid metabolism | 0 | 37 | 20 | 1 | 59 | 0 |

**Table S6**: Number of reproducibly identified proteins in the proteomes extracted by direct cell lysis using urea, sodium deoxycholate (SDC) and sodium dodecyl sulfate (SDS). Urea+SDC+SDS: number of identified proteins identified in both urea, SDC and SDS. Urea+SDC: number of identified proteins identified in both urea and SDC. Urea+SDS: number of identified proteins identified in both urea and SDS. SDC+SDS: number of identified proteins identified in both SDC and SDS. Urea: number of proteins identified in urea only. SDC: number of proteins identified in SDC only. SDS: number of proteins identified in SDS only.

| Condition | Number of identified protein groups | [%] |
|---|---|---|
| SDS+SDC+Urea | 4646 | 82.8 |
| SDC+Urea | 423 | 7.5 |
| Urea+SDS | 58 | 1.0 |
| SDC+SDS | 80 | 1.4 |
| Urea | 113 | 2.0 |
| SDC | 241 | 4.3 |
| SDS | 48 | 0.9 |

**Table S7:** Number of proteins extracted from SPM-LLE interphase pellets and by direct cell lysis with at least 1.5-fold higher efficiency.

| | **Urea vs. SDC** | |
|---|---|---|
| | # proteins up in urea | # proteins up in SDC |
| Direct cell lysis | 357 | 632 |
| SPM-LLE interphase pellets | 488 | 167 |

### Supplemental Experimental Section

#### Chemicals

HPLC-grade acetonitrile (ACN), methanol (MeOH), formic acid (FA), as well as Micro BCA™ Protein Assay Kit, Gibco® Qualified FBS and ammonium bicarbonate were obtained from Thermo Fisher Scientific (Dreieich, Germany). Dithiothreitol, iodoacetamide, HPLC-grade chloroform (CHCl3), urea, sodium deoxycholate (SDC), triethylammonium bicarbonate (TEAB), ethylenediaminetetraacetic acid (EDTA), Sera-MagTM magnetic carboxylate modified hydrophilic and hydrophobic beads were obtained from Merck and Sigma-Aldrich (Munich, Germany). Dulbecco's Modified Eagle Medium (DMEM), sodium dodecyl sulfate (SDS), sequencing grade modified trypsin, and HLB 1cc (30 mg) extraction cartridges, were purchased from PAN Biotech (Aidenbach, Germany), Carl Roth (Karlsruhe, Germany), Promega (Walldorf, Germany), and Waters Oasis (Vienna, Austria), respectively. [U-$^{13}$C]-labelled yeast extract was purchased from ISOtopic Solutions (Vienna, Austria). 1,2-Dimyristoyl-sn-glycero-3-phosphocholine (DMPC) and 1,2-dimyristoyl-sn-glycero-3-phosphoethanolamine (DMPE) were obtained from Avanti Polar Lipids (Alabaster, AL, USA), and 1,2,3-tri-myristoyl-glycerol (TMG) was purchased from EDQM (Strasbourg, France).

[U-$^{13}$C]-labelled yeast extract of Pichia pastoris (2 billion cells, ISOtopic solutions, Vienna, Austria) was reconstituted in 2 mL HPLC-$H_20$ aliquoted and stored at -80°C. 1,2- Dimyristoyl-*sn*-glycero-3-phosphocholine (DMPC, dissolved in $CHCl_3$, Avanti Polar Lipids, Alabaster, AL, USA), 1,2- dimyristoyl-*sn*-glycero-3-phosphoethanolamine (DMPE, dissolved in $CHCl_3$:MeOH 65:35, v/v; Avanti Polar Lipids, Alabaster, AL, USA) and 1,2,3-tri-myristoyl-glycerol (TMG, dissolved in $CHCl_3$, EDQM, Strasbourg, France) were combined and evaporated to dryness. The lipid film was taken up in a mixture of $CHCl_3$:MeOH:$H_2O$ (65:35:8, v/v/v) leading to the final concentration of 0.2 mM for each standard. The lipid standard was aliquoted and stored under argon at -80°C. All experiments were performed using five independent experiments (biological replicates).

#### Protein extraction from SPM-LLE interphase pellet using sodium dodecyl sulfate (SDS) and tryptic digestion

100 µg of total protein was used for tryptic digestion using single-pot solid-phase-enhanced sample preparation (SP3) procedure [1]. For this purpose, 50 µL each of carboxylate-modified hydrophilic and hydrophobic beads (Sera-Mag™, Merck, Sigma-Aldrich, Munich, Germany) were combined to a final concentration of 10µg/µL. Reaction vials containing the beads were placed on a magnetic rack for 2 minutes after which the supernatant was removed. 100 µL HPLC-$H_20$ were added to the beads and incubated for 30 seconds at RT to wash the beads. Reaction vials were incubated on a magnetic rack for 2 minutes, supernatant was removed and beads were suspended in 100 µL HPLC-$H_20$. For each sample, 100 µL of beads (total amount of beads: 1mg) was added to 100 µg of proteins dissolved in 100 µL lysis buffer (1 % SDS, 100 mM ABC, pH 8.3).) to reach a beads/protein ratio of 10:1 (wt/wt). 467 µL of acetonitrile (ACN) was added to the samples to reach a final concentration of 70% ACN. The samples were incubated for 18 minutes at RT, followed by an incubation on a magnetic rack for 2 minutes. Afterwards, the supernatants were transferred into new reaction vials, and the remaining reaction vials containing the beads were kept for further sample preparation (bead fraction 1). 100µL of a new mixed bead solution (total amount of beads: 1 mg) was added to the reaction vials containing the supernatants. The samples were incubated for 18 minutes at RT, followed by an incubation on the magnetic rack for 2 minutes. Supernatants were removed and the reaction vials containing the beads were kept (bead fraction 2). Both bead containing reaction vials (bead fraction 1, bead fraction 2) were washed twice by adding 200 µL of 70% ethanol, incubation at RT for 30 seconds and incubation on a magnetic rack for 2 minutes. The supernatants were removed. Additional two washing steps were performed by adding 300 µL 100% ACN incubation at RT for 30 s and incubation on a magnetic rack for 2 minutes. After removing the supernatant from the last washing step, beads were resuspended in 50 µL 50 mM ABC (dissolved in HPLC-$H_20$, pH 8.3). For reduction, 0.5 µL 1 M dithiothreitol (DTT, dissolved in 100 mM TEAB, pH 8.3) was added to the samples followed by an

incubation at 56°C for 30 minutes. For alkylation, 2µL 0.5 M iodoacetamide (IAA, dissolved in 100 mM TEAB, pH 8.3) was added to the sample followed and incubated for 30 minutes in the dark, followed by addition of 0.6 µL of reduction buffer to quench the alkylation reaction. Five µg of trypsin (dissolved in trypsin resuspension buffer, Promega, Walldorf, Germany) were added and the samples were incubated for 16 hours at 37°C. Digestion was stopped by addition of 1435 µL of 100% ACN to reach a final ACN concentration of 95%. Samples were vortexed and incubated for 8 minutes at RT. The supernatants were removed and the samples were washed twice with 200 µL ACN following the washing procedure described above. Finally, peptides were eluted from the beads by addition of 100 µL elution buffer (2% DMSO,1% FA, dissolved in HPLC-$H_20$). The samples were incubated on a magnetic rack for 2 minutes and the supernatants containing the eluted peptides from both bead fractions were combined to one sample. The supernatants were used for reversed phase solid phase extraction.

**Reversed phase solid phase extraction (RP-SPE)**

Samples were purified by RP-SPE prior to LC-MS analysis using OASIS HLB cartridges (Oasis HLB, 1 cc Vac Cartridge, 30 mg Sorbent, Waters, Manchester, UK) and a pressure manifold (Waters SPE Manifold, Waters, Manchester, UK). SPE cartridges were activated with each 1 mL of 100% methanol (MeOH), followed by 1 mL of 95 % ACN, 1% FA, and equilibrated with 1 mL of 1% FA. The samples were adjusted to a final volume of 1 mL and a final concentration of 1% FA. Samples were loaded on SPE cartridges and washed twice with 1 mL 1% FA. Peptides were eluted with 1 mL of 70% ACN, 1% FA. The solvents of the eluates were evaporated and dried peptide samples were stored at -80°C.

**Proteome analysis by LC-MS/MS**

Dried peptide samples were dissolved in 80 µL of 0.1% FA, and 1 µL of the samples were injected into a nano-ultra pressure liquid chromatography system (Dionex UltiMate 3000 RSLCnano pro flow, Thermo Scientific, Bremen, Germany) coupled via electrospray-ionization (ESI) to a tribrid orbitrap mass spectrometer (Orbitrap Fusion Lumos, Thermo Scientific, San Jose, CA, USA). The samples were loaded (15 µL/min) on a trapping column (nanoE MZ Sym C18, 5 µm, 180 µm x 20mm, Waters, Germany, buffer A: 0.1 % FA in HPLC-H2O; buffer B: 80 % ACN, 0.1 % FA in HPLC-H2O) with 5% buffer B. After sample loading the trapping column was washed for 2 min with 5% buffer B (15 µL/min) and the peptides were eluted (250 nL/min) onto the separation column (nanoEase MZ PST CSH, 130 A, C18 1.7 µm, 75 µm x 250mm, Waters, Germany; buffer A: 0.1 % FA in HPLC-H2O; buffer B: 80 % ACN, 0.1 % FA in HPLC-H2O). The peptides were separated using a total gradient of 110 min. First, peptides were separated using a gradient from 5% B to 37.5% B in 90 min, followed by 37.5% B to 62.5% B in 25 min. The spray was generated from a steel emitter (Fisher Scientific, Germany) at a capillary voltage of 1850 V. MS/MS measurements were carried out in data dependent acquisition mode (DDA) using an HCD collision energy of 30 % and top-speed scan mode. Every second a MS scan was performed over an m/z range from 350-1600, with a resolution of 120,000 FWHM at m/z 200 (maximum injection time= 50 ms, AGC target= 4e5, internal calibration mode activated using ETD reagent for mass calibration). MS/MS spectra were recorded in the ion trap (rapid scan mode, maximum injection time= 50 ms, AGC target= 1e4, quadrupole isolation width: 0.8 Da, intensity threshold: 1e4). Precursors were excluded from DDA analysis for 60 seconds.

**Bioinformatics data processing of proteome LC-MS/MS data**

The post-processing of the data was performed in R (version 4.0.3) and RStudio (version 1.4.1106). Statistical analysis to compare protein yields and number of identified protein groups was done by two-sided *t*-tests using rstatix package (https://cran.r-project.org/web/packages/rstatix/index.html). Eulerr package (https://cran.r-project.org/web/packages/eulerr/index.html, {Larsson, 2018 #17 was used to generate venn diagrams. For differential proteome analyses, the MaxQuant output file "proteingroups.txt" was used, which contained the quantified LFQ-values for the identified protein groups. LFQ values were log2-transformed and normalized to the median for each sample (normalized protein group intensities). For principal component analysis (PCA), the normalized protein group intensities of the proteins reproducibly quantified in all samples were used as an input. For volcano plots, two-tailed *t*-tests were performed for all protein groups and adjusted *p*-values were calculated using the Benjamini-Hochberg procedure using the rstatix package (https://cran.r-project.org/web/packages/rstatix/index.html). The r package "Peptides" (https://cran.r-project.org/web/packages/Peptides/index.html, {Osorio, 2015 #18} was used to calculate theoretical physicochemical properties of protein groups and GO enrichment analysis was performed using the gprofiler2 package (https://cran.r-project.org/web/packages/gprofiler2/index.html). The r ggplot2 package was used for data visualization (https://cran.r-project.org/web/packages/ggplot2/index.html) [2].

**Analysis of polar metabolites and lipids by Ion Chromatography-Single Ion Monitoring-Mass Spectrometry (IC-SIM-MS) and multiple reaction monitoring mass spectrometry (LC-MRM-MS)**

**Polar metabolites:**

The dried samples of the intracellular and extracellular methanol extracts were dissolved in 100 µL HPLC-$H_20$ and further diluted either 1:50 with HPLC-$H_20$ for the analysis of low abundant metabolites or 1:2000 for the analysis of high abundant metabolites. 4 µL of each sample were injected into a high-performance ion chromatography (HPIC) system (Dionex ICS-6000, Thermo Scientific, Germering, Germany). The separation was conducted on a Dionex IonPac AS11-HC column (2 mm × 250 mm, 4 µm particle size, Thermo Scientific) equipped with a Dionex IonPac AG11-HC guard column (Thermo Scientifc) at 35°C. A potassium hydroxide (KOH) gradient was produced by an eluent generator with a KOH cartridge (Dionex EGC 500 KOH, Thermo Scientifc) that was supplied with HPLC-$H_20$. For the separation, a flow rate of 380 µL/min and the following gradient were used: 0 mM KOH to 3 mM KOH in 3 min, 3 mM KOH to 10 mM KOH in 2 min, 10 mM KOH to 30 mM KOH in 15 min, 30 mM KOH to 50 mM KOH in 7 min, 50 mM to 85 mM KOH in 2 min. A Dionex AERS 500 suppressor was used to exchange potassium ions against protons in order to produce $H_20$ instead of KOH. A make-up flow (MeOH, 2 mM actic acid) was provided at a flow rate of 60 µL/min. A T-piece connected the IC-eluate and make-up flow with a heated electrospray ion source (HESI) of an orbitrap HF-X mass spectrometer (Thermo Scientific, Bremen, Germany). The following HESI source parameters were used: HESI temperature: 400°C, sheath gas: 50, auxiliary gas: 10, auxiliary gas temperature: 380°C, spray voltage: 2,500 V, S-Lens RF: 40, ion transfer capillary temperature: 380°C. MS analyses were performed in negative ion mode. Full MS spectra were recorded with a m/z scan-range of 80-520 m/z with a resolution of 60,000 FWHM at m/z 200, maximum injection time of 50 ms and an AGC target of 1e5. Metabolite quantification was carried out in targeted single ion monitoring (SIM) mode. Targeted SIM was acquired with a resolution of 60,000 FWHM at m/z 200, maximum injection time of 118 ms, AGC target of 1e5, an isolation window 4 m/z centered around the targeted m/z. The following targeted SIM windows were used:

**Table S8: List of polar metabolites and [U-13C]-labeled standards, retention times, m/z and charge [z] used for IC-SIM-MS.**

| Mass [m/z] | Charge [z] | Start t [min] | End t [min] | Metabolite |
|---|---|---|---|---|
| 179.05611 | -1 | 1.0 | 3.5 | Glucose |
| 87.00877 | -1 | 2.5 | 6.0 | Pyruvate |
| 259.02244 | -1 | 7.0 | 10.0 | Glucose-1-Phosphate |
| 117.01933 | -1 | 9.0 | 12.5 | Succinate |
| 133.01425 | -1 | 9.5 | 12.0 | Malate |
| 259.02244 | -1 | 10.0 | 16.0 | Glucose-6-Phosphate, Fructose-6-Phosphate* |
| 145.01425 | -1 | 12.8 | 15.0 | α-Ketoglutarate |
| 115.00368 | -1 | 14.1 | 16.0 | Fumarate |
| 289.03301 | -1 | 16.7 | 18.7 | Sedoheptulose-7-Phosphate |
| 346.05581 | -1 | 15.0 | 19.5 | Adenosine monophosphate |

| | | | | |
|---|---|---|---|---|
| 275.01736 | -1 | 22.0 | 24.0 | 6-Phosphogluconate |
| 173.00916 | -1 | 27.5 | 29.5 | Aconitic acid |
| 166.9751 | -1 | 27.5 | 29.5 | Phosphoenolpyruvate |
| 191.01973 | -1 | 25.4 | 29.0 | Citrate, Isocitrate* |
| 426.02214 | -1 | 30.5 | 32.0 | Adenosine diphosphate (ADP) |
| 337.98095 | -1 | 28.5 | 32.5 | Fructose-1,6-Bisphosphate |
| 505.98847 | -1 | 32.2 | 34.0 | Adenosine triphosphate (ATP) |
| 185.07624 | -1 | 1.0 | 3.5 | [U-13C]-Glucose |
| 90.01883 | -1 | 2.5 | 6.0 | [U-13C]-Pyruvate |
| 265.04257 | -1 | 7.0 | 10.0 | [U-13C]-Glucose-1-Phosphate |
| 137.02767 | -1 | 9.5 | 12 | [U-13C]-Malate |
| 265.04257 | -1 | 10 | 16 | [U-13C]-Glucose-6-Phosphate |
| 150.03102 | -1 | 12.8 | 15 | [U-13C]-α-Ketoglutarate |
| 119.0171 | -1 | 14.1 | 16 | [U-13C]-Fumarate |
| 356.08936 | -1 | 15 | 19.5 | [U-13C]-AMP |
| 281.03749 | -1 | 22 | 24 | [U-13C]-6-Phosphogluconate |
| 169.98516 | -1 | 27.5 | 29.5 | [U-13C]-Phosphoenolpyruvate |
| 197.03986 | -1 | 25.4 | 29 | [U-13C]-Citrate |
| 436.05569 | -1 | 30.5 | 32 | [U-13C]-ADP |
| 344.00108 | -1 | 28.5 | 32.5 | [U-13C]-Fructose-1,6-Bisphosphate |
| 467.02945 | -1 | 32.2 | 34 | [U-13C]-ATP |

* Both isobaric metabolites were measured in the same SIM window, as they exhibited baseline separation in IC.

Metabolite quantification was carried out with TraceFinder 5.0 General Quan (Thermo Scientific, San Jose, CA, USA). For peak identification and integration, the Genesis algorithm was used with the following parameters: Peak Detection Strategy: Highest Peak; Peak Threshold type: Area; Threshold: 1; Smoothing: 3; S/N threshold: 3; Tailing Factor: 3. If necessary, peaks were adjusted manually. Data were further processed using R (version 4.0.3) and RStudio (version 1.4.1106). R package rstatix (https://cran.r-project.org/web/packages/rstatix/index.html) was used to calculate the mean and standard deviation of the peak areas of the metabolites. The coefficient of variation (CV) was calculated by dividing the standard deviation by the mean and normalizing by a factor of 100. Violin plots of CVs were generated with R package ggplot2 (https://cran.r-project.org/web/packages/ggplot2/index.html) [2] and heatmaps were generated with R package gplots (https://cran.r-project.org/web/packages/gplots/index.html).

**Lipids:**

After SPM-LLE, lipid films were dissolved in MeOH (100 µl), centrifuged (21,100 x g, 4°C, 5 minutes), diluted 1:20 with MeOH and subjected to LC-MRM-MS after an additional centrifugation step (21,100 x g, 4°C, 5 minutes). Chromatographic separation of phospholipids (phosphatidylcholines (PC), lysophosphatidylethanolamine (LPE), and phosphatidylinositols (PI)) was performed using an ExionLC AD UHPLC system (Sciex, Framingham, MA, USA) as previously described [3]. Briefly, samples (injection volume: 3 µl) were separated at 45°C on an ACQUITY UPLC BEH C8 column (130Å, 1.7 µm, 2.1 × 100 mm; Waters, Milford, MA, USA) at a flow rate of 0.75 ml/min using buffer A (95% ACN, 2 mM ammonium acetate in $H_2O$) and buffer B (10% ACN, 2 mM ammonium acetate in $H_2O$). Lipids were separated with a gradient from 75% buffer A to 85% buffer in 5 minutes, followed by an increase to 100% buffer A within 2 minutes and a subsequent isocratic elution for another 2 minutes. Applying the instrumental set-up described above, triglycerides (TAG) were separated according to Espada et al. [4] at 45°C and a flow rate of 0.75 ml/min using a gradient consisting of buffer A (95% ACN, 2 mM ammonium acetate in $H_2O$) and buffer B (isopropanol). In short, the initial composition (90% buffer A) was reduced from 90% to 70%

within 6 min, which was succeeded by isocratic elution for 4 min. Eluted lipids were ionized by electrospray ionization (PC, LPE, PI: negative ion mode; TAG: positive ion mode) using a Turbo V ion source (Sciex, Framingham, MA, USA). Phospholipids were detected by MRM using a QTRAP 6500$^+$ Mass Spectrometer (Sciex, Framingham, MA, USA) following fragmentation of $[M+OAc]^-$ (PC) or $[M-H]^-$ ions (LPE, PI) to fatty acid anions, as described before [3]. For the simultaneous detection of PC, PE and PI, the curtain gas was set to 40 psi, the collision gas to medium, the ion spray voltage to -4500 V, the temperature to 500°C, the sheath gas to 55 psi, and the auxiliary gas to 75 psi. The declustering potential was set to -44 V (PC) or -50 V (PE, PI), the entrance potential to -10 V, the collision energy to -38 eV (PE), -46 eV (PC) or -62 eV (PI), and the collision cell exit potential to -11 V (PC, PI) or -12 V (LPE). TAGs were detected by a QTRAP 6500$^+$ Mass Spectrometer (Sciex, Framingham, MA, USA) in MRM mode by fragmentation of $[M+NH_4]^+$ adduct to [M-fatty acid acyl]$^+$ ions, without discriminating between fatty acyl positional isomers [3 4]. In variation to the settings described in the references above, the curtain gas was set to 40 psi, the collision gas to low, the ion spray voltage to 5500 V, the heated capillary temperature to 400°C, the sheath gas pressure to 60 psi, the auxiliary gas pressure to 70 psi, the declustering potential to 120 V, the entrance potential to 10 V, the collision energy to 35 eV, and the collision cell exit potential to 26 V. For quantification, the average of both transitions (PC, PE, PI) or the (most intensive) species-specific transition (TAG, lysophospholipids) was used and normalized to the internal standard (DMPC for PC, PI; DMPE for LPE; TMG for TAG) as well as the protein concentration [5]. The coefficient of variation (CV) was calculated using absolute intensities (analytes) or blank subtracted peak areas (DMPC, DMPE, TMG). Violin plots of CVs were generated with R package ggplot2 (https://cran.r-project.org/web/packages/ggplot2/index.html) [2] heatmaps were generated with R package gplots (https://cran.r-project.org/web/packages/gplots/index.html). The system was operated by Analyst 1.7.1 (Sciex, Framingham, MA, USA) and the obtained chromatograms were processed by Analyst 1.6.3 (Sciex, Framingham, MA, USA).
